# Generalized design of sequence-ensemble-function relationships for intrinsically disordered proteins

**DOI:** 10.1101/2024.10.10.617695

**Authors:** Ryan Krueger, Michael P. Brenner, Krishna Shrinivas

## Abstract

The design of folded proteins has advanced significantly in recent years. However, many proteins and protein regions are intrinsically disordered (IDPs) and lack a stable fold i.e., the sequence of an IDP encodes a vast ensemble of spatial conformations that specify its biological function. This conformational plasticity and heterogeneity makes IDP design challenging. Here, we introduce a computational framework for de novo design of IDPs through rational and efficient inversion of molecular simulations that approximate the underlying sequence to ensemble relationship. We highlight the versatility of this approach by designing IDPs with diverse properties and arbitrary sequence constraints. These include IDPs with target ensemble dimensions, loops and linkers, highly sensitive sensors of physicochemical stimuli, and binders to target disordered substrates with distinct conformational biases. Overall, our method provides a general framework for designing sequence-ensemble-function relationships of biological macromolecules.

## Introduction

The basis of biomolecular function is often specified by a sequence which encodes an ensemble of 3D conformations (1). A prominent example is intrinsically disordered protein regions (IDPs), which are found in most living organisms and play key roles in diverse cellular functions including transcription, cell signaling, cellular immunity, and translation (2–4). IDPs lack a stable 3D structure, rather, they dynamically interconvert between a large range of non-random conformations (5–7) whose local and global properties shape cellular functions (2). IDPs facilitate molecular recognition through embedded short linear motifs (8) and fuzzy interactions with multiple targets (2), and when tethered as intervening linkers or spacers, they modulate interactions between adjacent folded-domains (9). The conformational plasticity that underlies IDPs is highly sensitive to physicochemical and environmental contexts and thus they often function as intracellular sensors (10). Further, IDPs regulate assembly of higher-order biomolecular assemblies and condensates (11–14), often through low-affinity multivalent interactions, that play central roles in cellular signaling and information processing. Finally, dysregulation of IDPs and IDP-dependent interactions are increasingly correlated with multiple pathological states (11, 15). Thus, there is widespread interest to design IDPs with tailored functions for a variety of roles in human health and industry.

Despite recent advances in protein structure design enabled by the protein data bank (PDB) and machine learning (16–19), these computational methods have had limited ability for designing disordered proteins. Structures of IDPs are not characterized by single stable folds, rather, they occupy a vast ensemble of dynamic configurations. Recent developments in coarse-grained molecular simulations have successfully predicted *ensemble* properties of IDPs (20–22). These simulations produce training data for approximate machine learning models that predict particular properties (5, 23) (e.g. radius of gyration and polymer exponents) and can be subsequently inverted for design (24). While each method has found success, using separate algorithms for the forward and inverse problems reduces accuracy and generalizability to different target properties and force field parameters. It would be far preferable to *directly* invert the molecular simulations that model the sequence-ensemble relationship.

In this paper, we introduce an algorithmic approach to design IDPs with tailored properties through inverting molecular simulations. Our framework uses gradientbased optimization on molecular simulations for designing sequences with arbitrary equilibrium properties, bridging machine learning technology with ideas from statistical physics. We employ this method to engineer IDP sequences for a wide range and complexity of ensemble dimensions, including highly optimized loops and linkers. Our framework naturally accommodates arbitrary sequence constraints, which we highlight through the design of sequence patterning variants with the same composition but distinct ensemble properties. We then construct IDP-based sensors that are sensitive to salt concentrations, temperature, and concentrations of modification-driving enzymes. Finally, we design candidate IDP binders for highly disordered biological and synthetic substrates. Of note, the accuracy of our predictions is limited by the accuracy of simulation parameters that describe IDP sequence-ensemble relationships; our contribution is to show how to find optimal sequences *given a potential*. Our proposed method, while generically potential-agnostic, will benefit from the continued iteration between force-field development and experiment. Overall, our paper outlines a flexible strategy for *de novo* IDP design that can be generalized to engineer sequence-ensemble-function relationships for diverse biopolymers.

## Results

### Model Formulation

Rational de novo design of IDPs requires two key ingredients: (1) a reasonably accurate “forward” model of the sequence-ensemble-function paradigm (Figure 1A) and (2) an algorithm to “invert” this through directed search of sequence space towards a desired functional property. Over the last few years, coarse-grained molecular simulations with custom pair potentials have made (5, 21, 25) dramatic improvements in predicting effective ensemble properties of IDPs. In this paper, we focus on molecular dynamics simulations using 1 AA=1 bead coarse-graining with the Mpipi-GG (see Methods and SI Note 1) (21, 23).

**Fig. 1.**
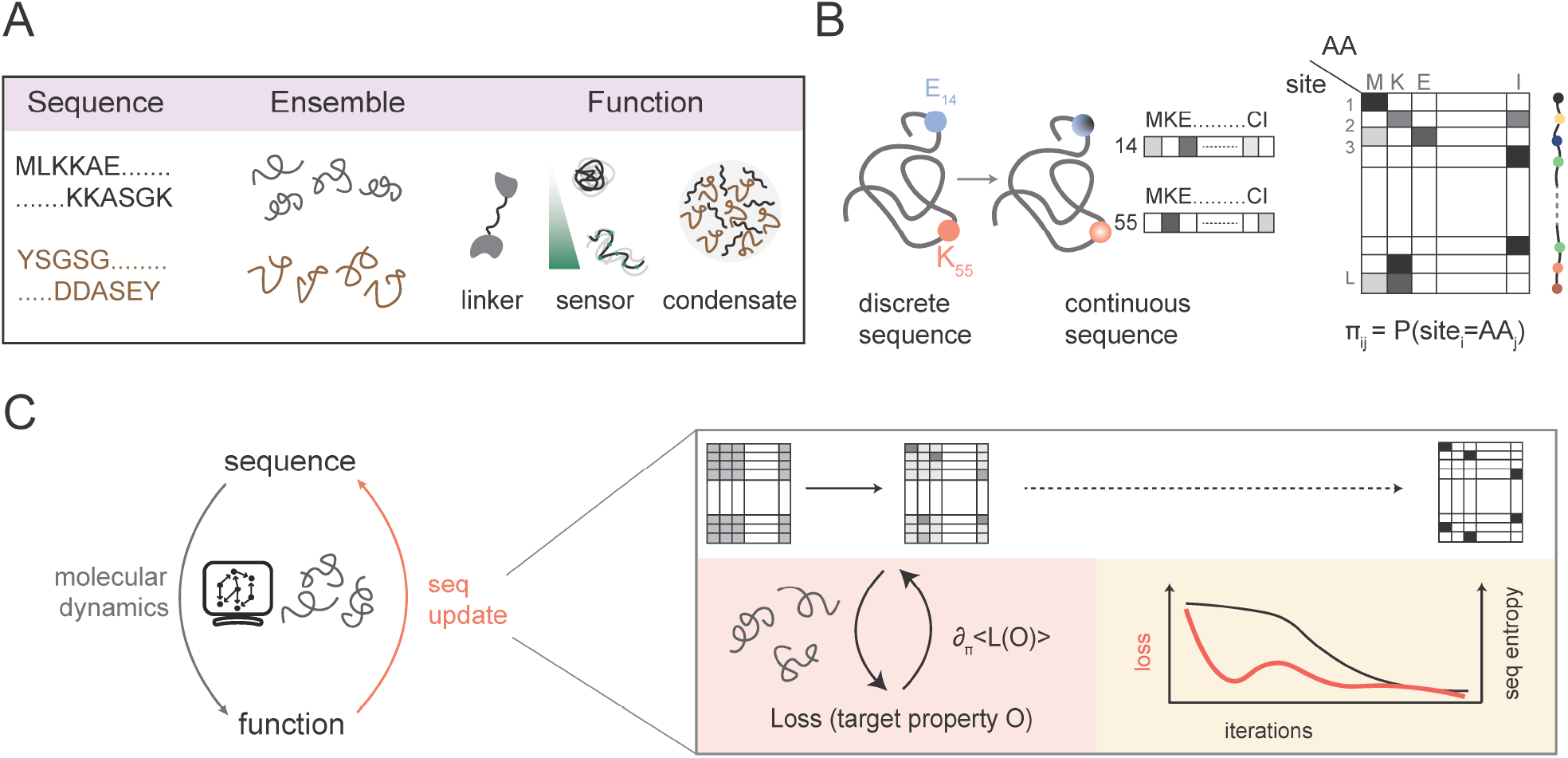
Method for inverse design of IDPs. **A**. The amino acid sequence of an IDP encodes for an ensemble of dynamic 3D conformations structures that determines properties shaping molecular and cellular functions. **B**. A discrete IDP sequence, a vector of length *n* where each position is typically a categorically represented amino acid character, is relaxed to a continuous, probabilistic sequence representation *π*, a matrix of size *n* × 20. Here, the (*i, j*) entry of *π* is the probability of residue at position *i* being amino acid *j*. **C**. To model the forward sequence-ensemble relationship, we simulate the probabilistic sequence through coarse-grained molecular dynamics simulations, defining the Hamiltonian of the system as the expected Hamiltonian over all sequences (see Methods). To invert this relationship for sequence design, we optimize this probabilistic sequence *π* via gradient descent and anneal to a discrete sequence through the optimization.

Our key innovation is the development of a differentiable algorithmic framework to *invert* the simulation-based sequence-ensemble relationships. To do this, we leverage recent advances in differentiable programming and stochastic gradient estimation (26–29) to compute the gradient of a loss function that depends on any set of ensemble-averaged properties: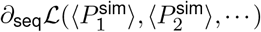. Since this quantity is only well-defined for smooth variable changes, we use a continuous representation of the sequence that is amenable to simulation and parallelization on GPUs (see Methods). For a sequence of *L* residues, this continuous probabilistic representation (Figure 1B), *π* = *f* (λ), is defined by logits λ of size *L* × 20. The residue identity at every site is characterized by a normalized probability vector over the different types of amino acids. A particular discrete sequence corresponds to a one-hot encoding i.e., each position is represented by a vector of length 20 with all entries but one being 0. In general, ensemble-averaged predictions are not identical to predictions from a distribution of discrete sequences sampled from the same distribution (see SI for derivation, Figure S2).

While in principle libraries like JAX-MD enable gradient calculation over unrolled MD trajectories, this is slow, scales poorly with system size, and is plagued by numerical instability (Figure S1, SI Note 2). To address this, we expand on a perturbative calculation developed independently by Zhang et al. (29) and Thaler and Zavadlav (28) to calculate the gradient with respect to *π* from a set of states sampled from the equilibrium Boltzmann distribution. This calculation provides significant speedup and accuracy in gradient estimation and allows reuse of simulation snapshots for multiple sequence updates. Finally, we incorporate an annealing procedure that gradually forces *π* to become increasingly discrete through the optimization (Figure 1C, see Methods). Unless otherwise specified, we initialize all optimizations with a uniform distribution.

### Designing IDPs with varying ensemble dimensions

Ensemble-averaged dimensions of an IDP, for e.g., the radius of gyration (*R*_*g*_) or the end-to-end radius (*R*_*ee*_), are coarse-grained metrics that reveal conformational biases which can correlate with binding and emergent phase behavior (20, 30, 31). Therefore, we first set out to design an IDP of fixed sequence length (*n* = 50) with a target dimension of ⟨*R*_*g*_⟩ = 20 Å. We then update *π* in the direction of desired ⟨*R*_*g*_⟩ while simultaneously annealing, albeit gradually, towards a discrete sequence (Figure 2C). Our routine converges (over 50 epochs and 2.5 hours on an NVIDIA A100 GPU) to a sequence (Figure 2B, Supplementary Data, SI Note 3) which explores a range of conformations (Figure S2) with an ensemble-averaged *R*_*g*_ of ∼ 20.1 Å. Rerunning the optimization with varying random seeds leads to different sequences with similar *R*_*g*_ – highlighting the ability of our approach to identify multiple sequences that exhibit similar ensemble-averaged properties (Figure S2, Supplementary Data).

**Fig. 2.**
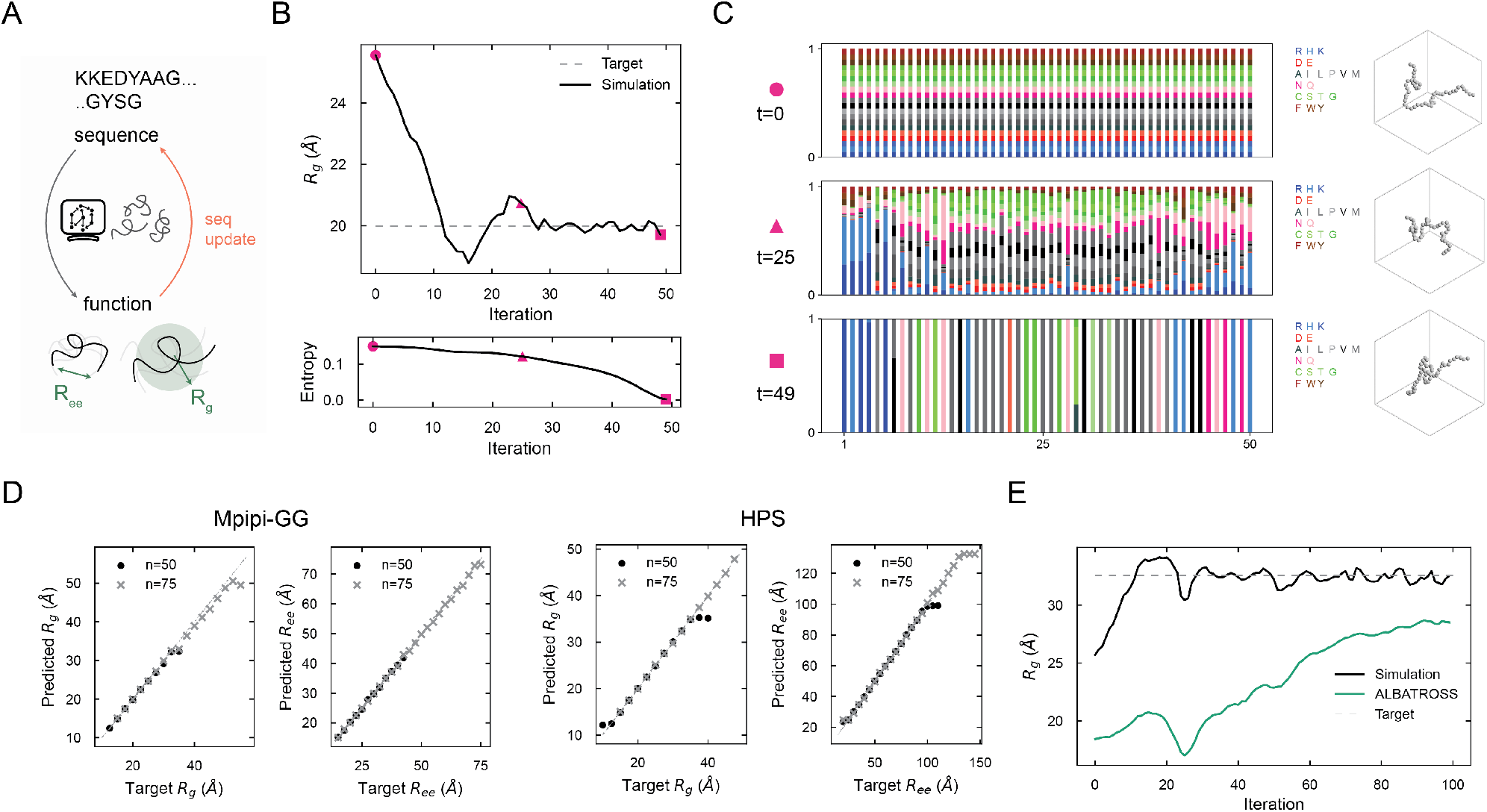
Designing IDPs with varying ensemble dimensions. **A**. Employing the framework defined in Figure 1 for design of IDPs with defined ensemble-averaged physical dimensions, specifically the *R*_*g*_, *R*_*ee*_. **B**. An example optimization to design an IDP of length *n* = 50 with *R*_*g*_ = 20 Å. The top panel represents the *R*_*g*_ from the simulated probabilistic sequence and the bottom panel represents the average sequence entropy at each position. Highlighted points (in pink) represent approximately the start, mid-point, and end of the optimization. **C**. The evolution of the probabilistic sequence throughout the optimization depicted in **B**. at highlighted points accompanied by a characteristic conformation in a box of side *a* = 75 Å. Each residue is colored differently and the column height corresponding to each residue position is the likelihood of being each residue. The probabilistic sequence is initialized as a uniform distribution of sequences, with each residue having an equal probability at each position, and the final sequence is nearly discrete. **D**. Each panel shows results for a set of optimizations, with each point comparing the predicted versus target ensemble dimension (*R*_*g*_ or *R*_*ee*_) for a particular IDP sequence. The different panels highlight solutions for different sequence lengths (*n* = 50, 75) and for different force-fields (Mpipi-GG - left two panels, HPS - right two panels). **E**. The optimization trajectory for a sequence of length *n* = 50 for target *R*_*g*_ = 35 Å in which ALBATROSS underpredicts *R*_*g*_ of the final optimized sequence by ∼ 4 Å.

With this framework, we are able to generate sequences of multiple lengths (*n* = 50, *n* = 75) across a wide span of *R*_*g*_ (Figure 2D). When we change the loss to correspond to a different physical property, the end-to-end radius or *R*_*ee*_ – a dimension which provides insights into linker function in multi-domain proteins (9) – we are able to design IDPs across a wide range of *R*_*ee*_ (Figure 2D, SI Note 4). We find that the optima we obtain using this method are more accurate than those obtained with a pure machine-learned predictor derived from Mpipi-GG simulations (ALBATROSS), when compared against the underlying molecular dynamics simulations for ground truth (Table S1). As an example, a sequence we generate (*n* = 50, ⟨*R*_*g*_⟩ = 32.55 Å, ⟨*R*_*g*_⟩ ^target^ = 32.5 Å) is incorrectly predicted by ALBATROSS to be off by ∼ 4 Å (Figure 2E). A core strength of our algorithm is that by directly optimizing over simulations, we can explore a wider design space that is not subject to approximations underlying machine-learned descriptors. This means that more generally, our method can be flexibly applied to any force field without requiring further data generation, architecture engineering, fine tuning, or retraining of existing models. We demonstrate this by designing IDPs of particular ensemble dimensions using the same method but with a different commonly used pair potential (Figure 2D). Together, our method provides a versatile approach to identify IDPs with specified conformation-averaged single-chain properties.

### De novo design of loops and linkers

We next asked, can we construct IDPs with more complex descriptors of their conformational ensembles? In particular, we focused on designing sequence variants that maximized decoupling between *R*_*g*_ and *R*_*ee*_ as opposed to the linear scaling found in ideal polymers, unfolded proteins, and many naturally occurring IDPs (32, 33). We reasoned that such sequence variants could potentially represent optimally designed loops 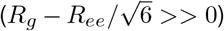 or linkers 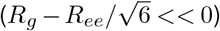 (Figure 3A, SI Note 5).

**Fig. 3.**
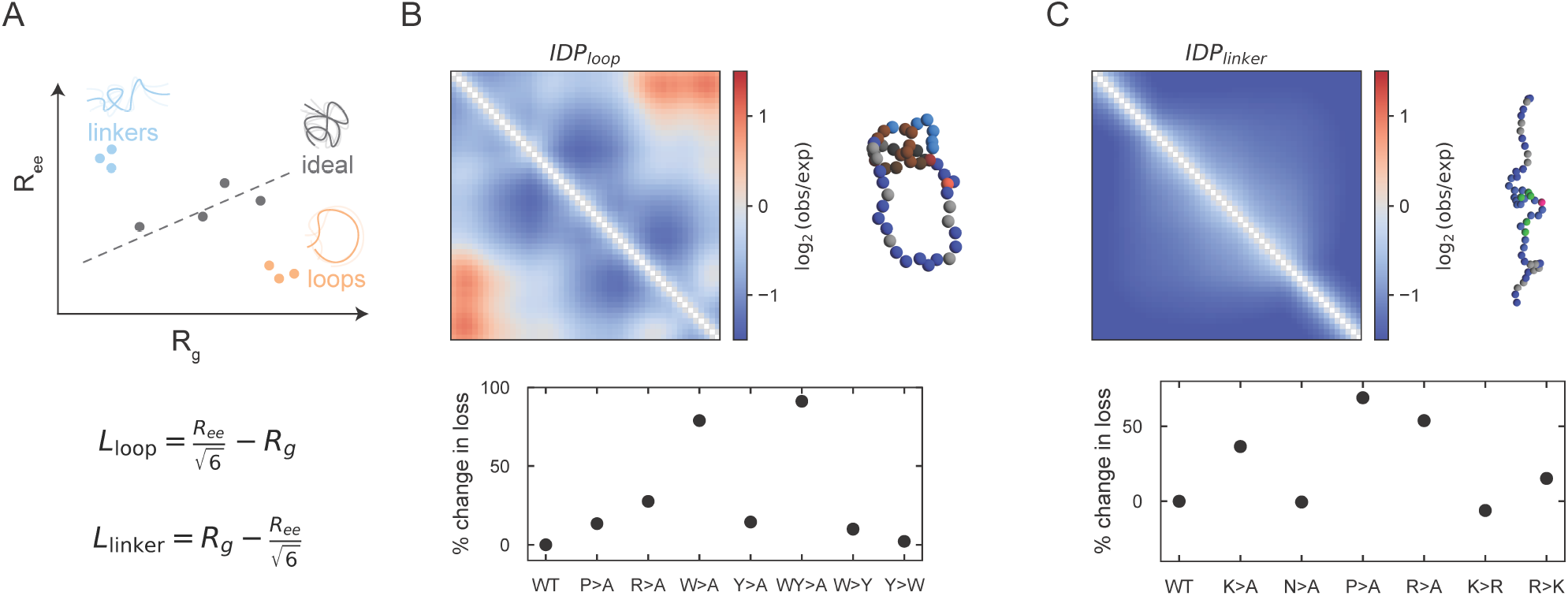
Shaping global conformational biases through loops and linker IDPs. **A**. A graphical illustration of ensemble coupling of *R*_*g*_ and *R*_*ee*_, highlighting linear relationships for ideal homopolymer chains 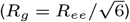 and decoupled off-diagonal points for loops and linkers. Below, we show the loss we employ for the loop and linker design problems, expressed to maximize the decoupling between *R*_*g*_ and 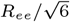 **B-C**. For the optimized loop (**B**) and linker (**C**) sequences, we depict the normalized contact frequencies computed over a trajectory. To the right of it, a representative configuration that is colored by amino acid identity (similar to Figure 2C), and below it, the relative change in loss value for a set of key mutational scans. For contact frequencies, red/blue regions represent higher/lower expected frequencies when contrasted with an ideal polymer of identical length. The generic increase in loss upon mutation represents that our solution is highly optimized for the target property.

For a sequence of fixed length (*L* = 50), we identify highly optimized loop and linker sequences with finely-tuned mechanistic properties. Our loop optimization yields a low-complexity sequence with sticky aromatic patches comprising tryptophans and tyrosines at either termini, interspersed by prolines and arginines that kink out the intervening sequence – highlighted by the normalized contact frequency maps and representative conformations (Figure 3B). Although the underlying force-field predicts that W-W interactions are stickier and perhaps should thus drive stronger loops, mutating the mixture of Y/Ws in our solution to either all Ys or Ws leads to a less optimal loop (Figure 3B, Supplementary Table S2). Similarly, mutational scans of each residue type into alanines or choosing less-complex losses lead to suboptimal loops (Supplementary Table S2) – generically reflecting an inability of simple sequence perturbations to decouple reductions in end-to-end distances from concomitant reductions in chain *R*_*g*_. Hence, the optimal loop architecture here arises from tradeoffs between overall sequence composition and patterning and emergent many-body interactions. When optimizing for linkers, we find that low-complexity sequences that intersperse prolines amongst a backbone of positively charged arginines, maximally decoupling *R*_*ee*_ from *R*_*g*_ (Figure 3C). This is largely expected since like-charges have short-range repulsive interactions and simple mutation scans (Figure 3C) are consistent with this intuition. Interestingly, we still identify a variant (R → K) that leads to slightly more optimal linkers. Overall, these design problems reinforce the ability of our algorithm to navigate high-dimensional sequence-spaces while balancing tradeoffs in ensemble properties.

### Engineering IDPs with arbitrary sequence constraints

An important aspect of protein design is to engineer molecules that are subject to sequence constraints. For IDPs, such constraints could span requirements for highly disordered sequences, particular sequence compositions or motifs, or any other combinatorial sequence features. To incorporate arbitrary constraints, we generically expand our algorithmic framework by building on our previous work (34). First, constraints are enforced through leaky ReLu functions multiplying the target property loss, resulting in gradients that navigate sequence space while maintaining constraints (Figure 4A). Second, instead of directly optimizing over the sequence, we optimize over the weights of a pre-trained and fully connected NN that parametrizes *π* (Figure 4A). Together, this presents a modular and generalizable strategy to navigate constrained high-dimensional sequence spaces (SI Note 6).

**Fig. 4.**
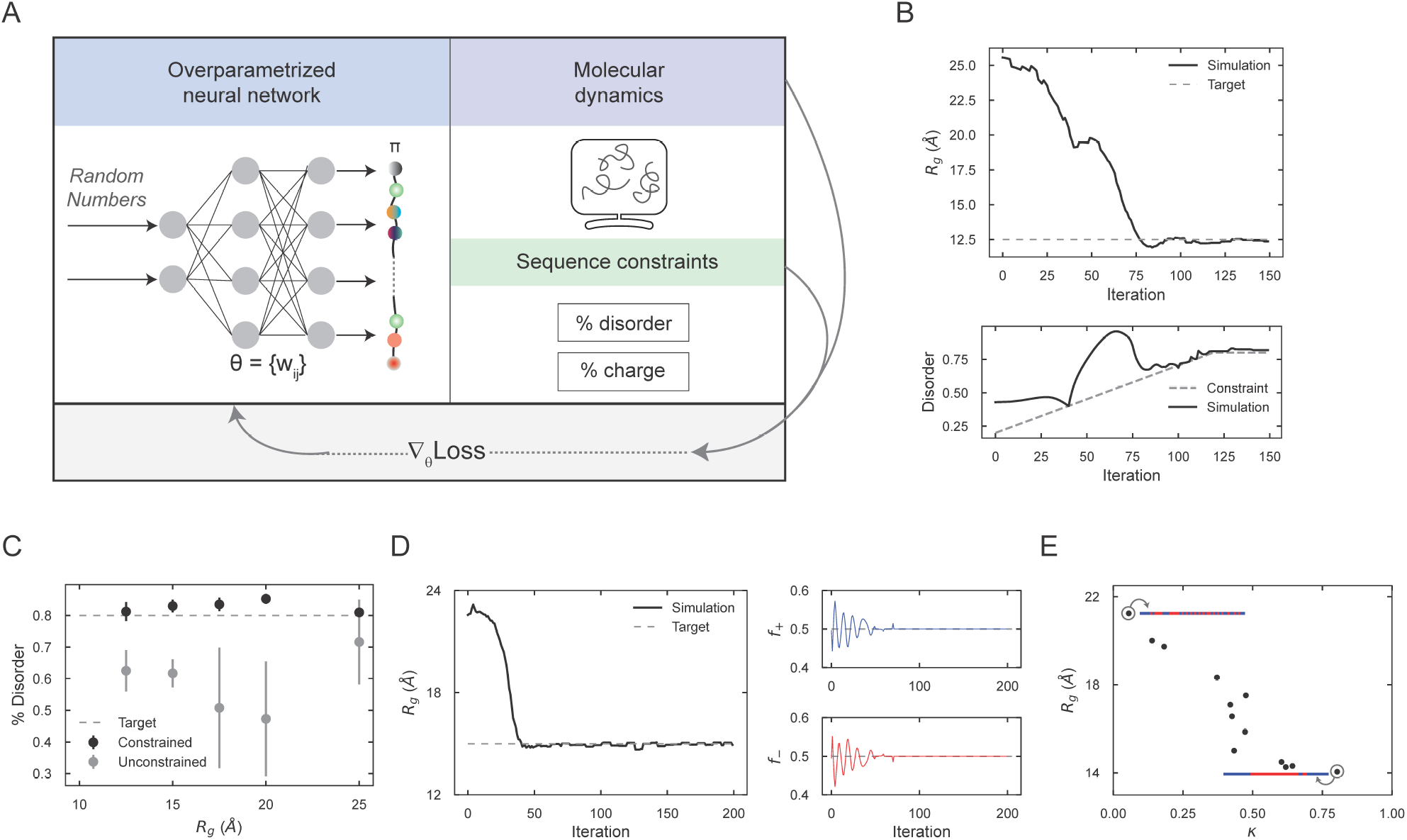
Engineering IDPs with arbitrary sequence constraints. **A**. Our framework for applying sequence constraints. Following Ref. (34), we construct a loss function that incorporates arbitrary constraints on the probabilistic sequence and overparameterize the input to the optimization problem i.e., the sequence representation, via a neural network. **B**. An example of IDP design (*n* = 50, *Rg* = 12.5 Å) subject to a constraint requiring a minimum degree of sequence disorder as predicted by Metapredict (35). The top panel shows the simulation-predicted *R*_*g*_ over training epochs. The bottom panel shows the annealing of the sequence disorder constraint across the optimization. **C**. Average disorder of optimized sequences (*n* = 50, 5 replicates) versus target *R*_*g*_ value with (black) and without disorder constraints (grey). Dashed lines represent the threshold of enforced disorder constraint. Optimized sequences exhibit a *R*_*g*_ within 5% / 10% of target value for constrained/unconstrained optimizations. **D**. An example of IDP design (*n* = 50, *R*_*g*_ = 17.5 Å) subject to a constraint that requires 50% positively charged (R/K) and 50% negatively charged (E/D) residues. The constraints (right panel) and the target Rg (left panel) are achieved as the probabilistic sequence is annealed to a discrete sequence. ***E***. *R*_*g*_ values of optimized sequences are plotted against *κ* values, a measure of sequence blockiness introduced in (36), recapitulate the inverse relationship shown in (36). The inset depicts sequence patterning represented by blue/red lines for positive/negative residues and shows relatively interspersed/blocky IDPs for high/low *R*_*g*_ values. Each dot represents the most optimized sequence from 5 trajectories and have an *R*_*g*_ within ∼ 10% of the target and charge ratios within ∼ 5% of target.

With this framework, we first set out to identify IDPs that are constrained to be highly disordered. We leverage a recent ML-based disorder predictor, Metapredict (35), to measure and constrain disorder (SI Note 6). Importantly, since the disorder prediction (and requirement) is only exact for a discrete sequence, the disorder-contribution to the loss is gradually made more stringent over the optimization procedure (Figure 4B). Designing compact proteins i.e., those with small *R*_*g*_, without any constraints tends to discover highly hydrophobic proteins that are typically predicted to be well-folded and not disordered (see Figure 4C). When we incorporate our disorder constraint, we are able to identify sequences that are simultaneously compact and highly disordered (Figure 4B) across a range of *R*_*g*_ (Figure 4C).

We next set out to design IDPs with *compositional* constraints. Motivated by previous work (36), we explored the effect of sequence patterning, particularly blockiness, on ensemble dimensions while keeping overall composition fixed at 50% positive and negative charges. To perform this multi-constraint optimization (Figure 4D), we pretrain the overparameterized fully-connected NN to output a set of logits corresponding to the target charge distribution, and then use this in our constrained optimization procedure. Consistent with previous predictions, we find an inverse relationship between ensemble dimensions and sequence blockiness (Figure 4E). Together, our results demonstrate the ability of our model to design IDPs with multiple sequence-based constraints.

### Programming stimuli-response in IDPs

A key biological function of many IDPs stems from their ability to sense and respond to cellular and environmental stimuli such as varying salt concentrations, temperature changes, dissolved CO_2_ levels, and pH (10, 37) through changing global or local chain conformations. Thus, we next decided to create IDP-based sensors, where we defined sensor function as arising from large changes in global conformation (*R*_*g*_) in response to varying external stimuli (Figure 5A, SI Note 7). Our algorithm naturally handles such complex design formulations, which require tailored sequence-ensemble-function relationships across multiple conditions: for example, a salt contractor IDP sensor must have high *R*_*g*_ at low salt and low *R*_*g*_ at high salt concentrations. Thus the design optimization must find the sequence that achieves this goal simultaneously over *both* conditions.

**Fig. 5.**
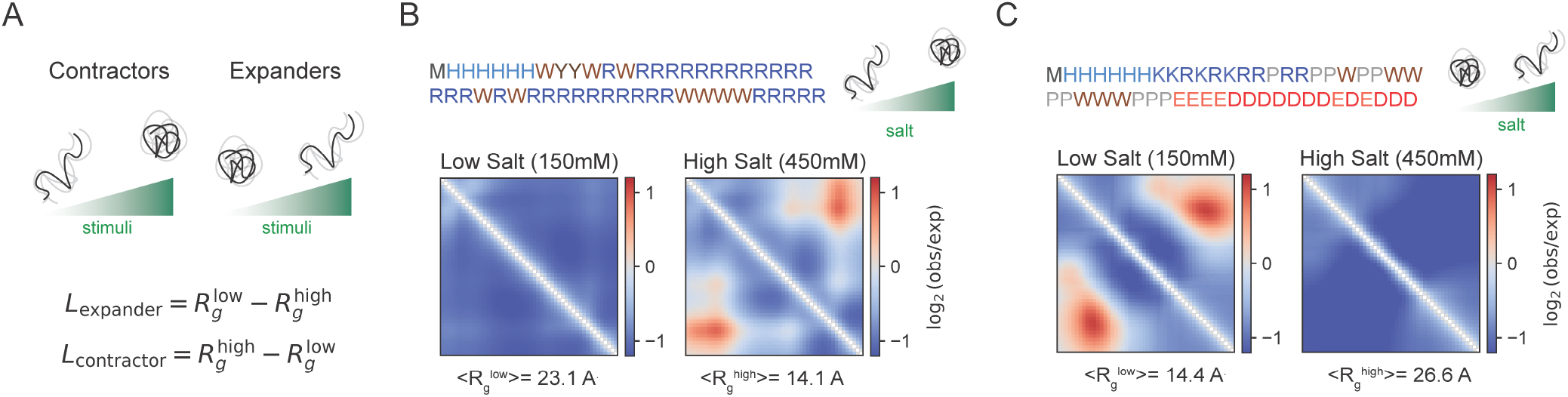
Programming stimuli-responsive IDP sensors. **A**. The stimuli-response is depicted by global change in ensemble dimensions across the stimulus (green gradient bar could mean salt, temperature, or phosphorylation) and contractors/expanders represent the direction of change. Below the illustration, we show the specific loss that is expressed as the difference of the two *R*_*g*_ values across the stimuli conditions, with the sign depending on whether we are designing a contractor or expander. **B-C**. Optimized contractor and expander sequences with contact frequency maps at low and high salt computed from representative trajectories and *R*_*g*_ is reported below. For contact frequencies, red/blue regions represent higher/lower expected frequencies when normalized with an ideal polymer of identical length.

We first began by designing sensors that respond to an increase in salt concentrations from 150 mM to 450 mM, where the salt concentration affects electrostatic screening lengths (SI Note 7). By optimizing for a salt-contractor, we identify a sequence (Figure 5B) rich in arginines with small clusters of interspersed H/Y/W residues. The weakening of repulsive interactions between like-charge R’s with salt leads to an effective and modest compaction – as exemplified by a poly-R sequence of identical length (Table S4). In our solution, this passive contraction is amplified by the aromatic clusters whose attractive bonding is salt-insensitive. The periodic spacing and patterning of aromatic solutions in our designed variant only drives compaction under high salt conditions (Figure 5B). All but one mutants that change composition or patterning lead to more compaction but are no longer as salt-sensitive (Table S4, Figure S4A, Supplementary Data).

This ability to exploit complex, many-body heteropolymer physics is even more dramatic in our salt-expander variant (Figure 5C), designed to *increase R*_*g*_ with increasing salt concentration. Our algorithm converges to an expander with 3 roughly equal-size sequence modules: positively charged N-terminus, negatively charged C-terminus, and a linker region that is made of proline spacers interspersed with sticky aromatic residues. The weakening of attractive interactions between positive and negative residues with salt only drives modest expansion, as seen in a K_25_E_25_ variant (Table S4, 2.2 Å change). Two linker features, (a) a sticky pi-cation interactions with aromatic residues and the N-terminus and (b) steric effects from proline residues that reduce contact frequency of N/C termini, work in tandem to drive a salt-sensitive “molecular-clasp” (Figure 5C) with a nearly 100% change in *R*_*g*_. Removal of any key features, e.g. through reducing steric hindrance by P → A mutations, or removing sticky residues, leads to a weaker salt-response (Figure S4B). Thus, our method identifies a balance between salt-sensitive, salt-independent, and steric features whose coupling transforms into a cooperative large-scale stimuli-response. This designed molecular clasp, in turn, sheds light on physical mechanisms that underlie sensitive and plastic conformational ensembles.

Finally, to highlight the generality of our model, we use a similar approach to construct sensors that respond by contraction or expansion to increases in temperature and to phosphorylation of serine residues (Figure S5, Supplementary Table S3, SI Note 7). Since the underlying force-fields do not accurately capture temperature dependent variations in hydrophobic interactions, the effect sizes predicted by our model are rather small (Figure S5D-F). Incorporating temperature dependent interactions e.g., in the spirit of Ref. (38), into our model framework will improve future sensor design. By contrast, we find an increasing range in sensor dimension change, and thus response size, with more phosphosites (Figure S5A-C). The mechanisms of contraction/expansion rely on interactions between phosphorylatable residues buried in neighborhoods of positively/negatively charged residues (Figure S5A-C). Thus, the addition of negatively charged groups upon phosphorylation promotes favorable or repulsive interactions, leading to downstream change in IDP ensemble size.

### Binders for disordered substrates

The function of many IDPs is driven by binding to disordered substrates, with examples of pico-molar level affinities in highly charged IDPs (30, 39, 40). We next asked, can we design disordered binders for a specific target substrate? To do this, we modify the forward simulation to include both the substrate, whose residue identity is fixed but can still sample a variety of conformations, along with a potential binding ligand whose sequence is learnable (Figure 6A). Precise calculation of binding constants is computationally expensive, often requiring sophisticated enhanced sampling techniques. To overcome this, we make the following simplifications: (a) strong binders are identified by minimizing average interstrand distance and (b) a biasing potential is employed to encourage collection of reference samples that are confined to an effective local volume (SI Note 8). These simplifications help identify high-affinity binders but lose the ability to measure precise quantitative rates or constants.

**Fig. 6.**
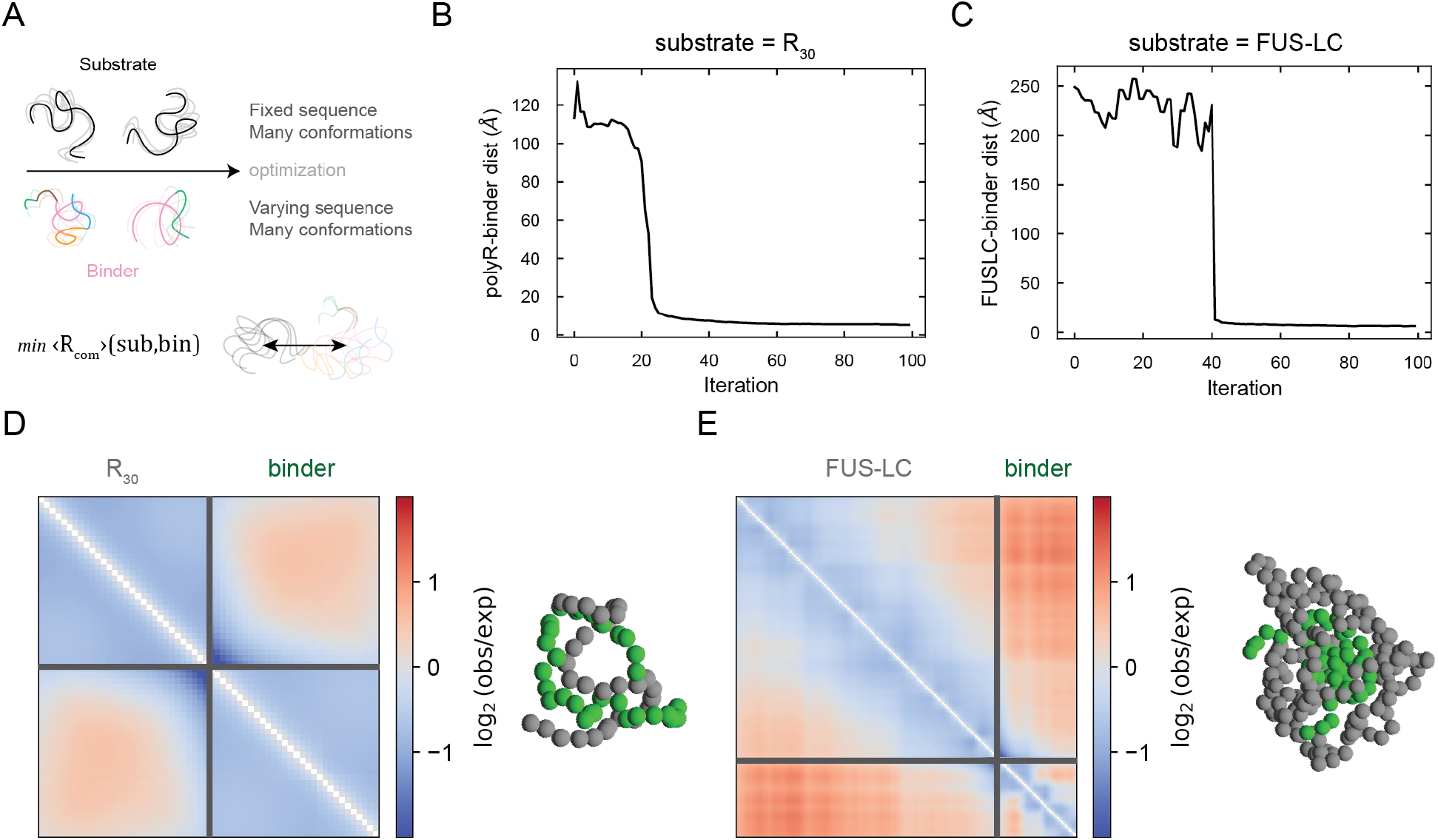
Designing IDP binders for disordered substrates. **A**. For a given substrate, binder design involves finding an IDP sequence that minimizes the interstrand center-of-mass distance between conformationally fluctuating IDPs. **B**. The optimization of a binder of length *n* = 30 for *R*_30_ is depicted by convergence to low interstrand distance over the trajectory. **C**. The optimization of a binder of length *n* = 50 for FUS-LC (*n* = 169) is depicted by convergence to low interstrand distance over the trajectory. **D-E**. Normalized contact frequency maps for *R*_30_ (D) and FUS-LC (E) are shown, highlighting both intramolecular and intermolecular interactions. For contact frequencies, red/blue regions represent higher/lower expected frequencies when contrasted with an ideal polymer of total length of binder + substrate. Representative bound snapshots of binder (green) and substrate (grey) are depicted to the right of the panel and lines within box separate out substrate and binder.

We first seek a binder for a homopolymeric positively charged substrate R_30_. Our model identifies a predominantly negatively charged ligand (>90% D/E residues, Supplementary Data) as a strong binder. Consistent with poly-electrolyte models, we find that predicted effective interaction coefficients (41, 42) (SI Note 8) are highly favorable for unlike-charge mediated substrate-ligand interactions (Figure S6C) and unfavorable, as expected, for like-charge mediated substrate-substrate interactions. We next identify binders for the Low-Complexity domain of FUS, a well-studied IDP with prominent roles in human physiology and disease (43, 44) and the poly-Q region of Whi3, an IDP with prominent roles in regulating nuclear autonomy and cell cycle in budding yeast (45). As shown in Figure 6B, our optimization leads to an identification of a target binder (Supplementary Data). While both FUS-LC and Whi3 have strong self-affinity (43, 45), effective interaction coefficients predict stronger interactions between our optimized binders and their respective substrates (Figure S6C) over the homotypic substrate-substrate interactions. In unbiased forward simulations, we observe strongly enriched intermolecular interactions for all binder-substrate pairs (Figures 6D-E, S6B), indicating strong binding at the *µM* concentrations we studied. Across all the optimizations, we find that a sharp change in the learning dynamics (Figures 6B-C, S6A) is concomitant with strong binder identification. We expect that future studies will dissect whether this transition represents features of the underlying learning protocol i.e., annealing schedule or noisy gradient signal due to limited sampling, or reflects the cooperative biophysics of such molecular binding events. Overall, our model lays the framework to generate candidate IDP binders for disordered substrates.

## Discussion

Intrinsically disordered proteins and protein regions (IDPs) are biomolecules that are found across the tree of life, play critical roles in molecular recognition, cellular organization, and information processing, and when dysregulated, correlate with pathology. The sequence of an IDP encodes for a vast repertoire of interconverting spatial conformations that shape their emergent function. *De novo* design of IDPs with diverse and arbitrary properties remains limiting, in large part, due to lack of methods to generally invert the underlying sequence-ensemble-function relationship.

In this paper, we introduce a computational framework to discover IDPs for a wide variety of target functions by rationally and efficiently inverting molecular simulations that capture the underlying sequence-ensemble relationship (Figure 1). Using this framework, we first design IDP sequences with varying and complex coarse-grained ensemble dimension properties. Specifically, we design sequences across a range of *R*_*g*_ and *R*_*ee*_ (Figure 2), and with tailored conformational biases i.e., loops and linkers (Figure 3), properties that have been shown to shape cellular function (2, 30). We next develop a modular strategy to incorporate any sequence constraints in the design pipeline. With this, we engineer IDPs that are simultaneously compact and disordered, and generate sequence patterning variants with the same overall composition but differing ensemble dimensions (Figure 4). With this framework, we next design highly sensitive sensors to multiple physicochemical and cellular stimuli such as salt, temperature, and phosphorylation (Figure 5). These designed sensors, in turn, shed light on the physical mechanisms by which the balance of competing intramolecular interactions encodes for large conformational changes. Finally, we use this method to identify disordered binders for low-complexity substrates (Figure 6). More generally, there is significant potential to apply the outlined framework to distinct biomolecular sequences (proteins, RNA, DNA) with equilibrium sequence-ensemble-property relationships that can be predicted by a wide range of techniques spanning molecular dynamics, Monte-Carlo simulations (46), field-theoretical approaches (47), and thermodynamics-informed models (41, 42, 48).

The framework we propose directly inverts simulation-derived sequence-ensemble relationships to drive *de novo* IDP design with tailored single-chain, binding, and environmental-specific properties. A key aspect of this approach is the integration of continuous and relaxed sequence representations with molecular simulations, inspired by a host of recent efforts that invert analytical calculations or machine-learned approximations for biopolymer design (34, 49, 50). Predictions via our framework, which require experimental tests, are fundamentally constrained by the accuracy of the underlying simulations. A key advantage of our approach is the underlying flexibility of gradient-based optimization, which in principle, can be leveraged to calculate gradients and optimize *simulation parameters* instead of sequence design. Namely these same methods can be used in combination with experiments to drive an iterative loop to improve simulation accuracy that is benchmarked on multimodal experimental measurements of IDP properties. In a parallel paper we demonstrate how such an approach can improve simulation accuracy by fitting the parameters of a coarse-grained model of DNA to complex experimental data such as melting temperatures and stretch and torsional moduli (51).

Incorporation of emerging machine-learning approaches – for e.g., simulation-free generative methods to generate conformational ensembles (52, 53), combining alchemical and molecular dynamics simulations for sequence variant design with target single-chain properties (5, 54), and approximate ML models that can rapidly invert pre-trained sequence-single-chain property relationships (23, 54), continues to expand the toolbox for protein engineering. Combining physics-based approaches with recent advances in differentiable programming holds promise for computational design and engineering for a wide variety of biomolecules and their functions.

## Limitations of the study

Our paper introduces a framework to design IDPs with tailored equilibrium sequence-ensemble relationships modeled by simulations. First, since we compute gradient estimates via a reweighting scheme that relies on knowledge of unnormalized probabilities, our framework in its present form does not naturally accommodate far-from-equilibrium properties for which state-level probabilities are generally not known. Opportunities to address this limitation in future work include exploiting classic results in non-equilibrium statistical mechanics (e.g. Jarczynski equality), jointly learning the parameters of the attractor of a dynamical system (similar to actor-critic methods in reinforcement learning), and alternative methods of automatic differentiation that sacrifice accuracy for numerical stability and memory overhead. Second, the convergence of this approach to niche sequence-designs has not been stress-tested and may require further algorithmic innovations. A particular challenge for convergence is the inequality between ensemble statistics computed via a continuous representation versus a distribution of discrete sequences sampled from the continuous sequence. Third, we only explored models for which the geometry of each particle identity is identical and probing models with polydisperse and complex geometries may require further methods development. Finally, directly inverting molecular simulations has a direct tradeoff contrasting increased accuracy with additional speed and compute requirements, thus making it less appealing for design of properties for which machine-learned approximations are comparable in accuracy.

## Methods

### General framework for optimizing particle identities

Consider a system of *n* particles in *d* dimensions where each particle is ascribed one of *m* possible identities. Let 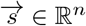 denote the identities of each particle where 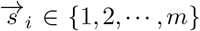 Given a potential energy function *U* : ℝ*n*×*d* → ℝ that depends on the particle identities, 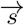 determines the distribution of states in the canonical ensemble via 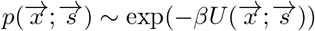 where *β* is the inverse temperature and 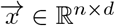. Given some statelevel observable *O* : ℝ*n*×*d* → ℝ, one is typically interested in the expected value of *O* in the entire ensemble,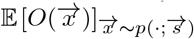. Consequently, we consider the optimization problem

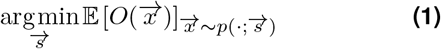

Note that this is equivalent to the maximization or fixed point variants of the optimization problem.

We define an optimization framework for Equation 1 that

(i) is general and makes minimal assumptions about the underlying model, (ii) operates directly at the level of the model and requires no training, (iii) yields an optimized probability distribution of identities from which discrete identities can be sampled, and (iv) can be combined naturally with state of the art machine learning methods. Consider a matrix of particle identities, *π* ∈ ℝ*n*×*m*, where *π*_*ij*_ is the probability of the *i*^*th*^ particle having identity *j* and Σ*π*_*ij*_ = 1.0 for all *i*. Let *S* denote the set of all possible discrete vectors of particle identities with |*S* |= *m*^*n*^. We can then define the expected potential energy of a state 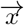 as

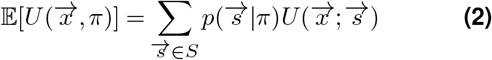

where

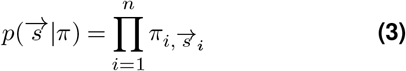

This yields a corresponding distribution of states in the canonical ensemble,

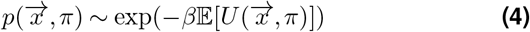

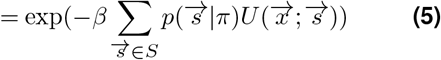

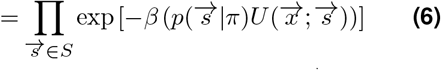

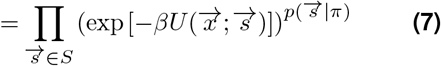

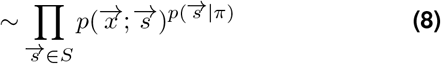

Given this generalized probability distribution, we can generalize Equation 1 for the case of probabilistic particle identities:

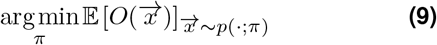

Note that Equation 9 reduces to Equation 1 in the case where *π* is one-hot.

Crucially, *π* is a continuous variable and can be optimized via gradient descent. Given a stochastic sampler (e.g. a Langevin integrator), one can compute 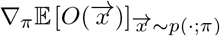 via Differentiable Trajectory Reweighting (DiffTRE) (28). Consider a set of states 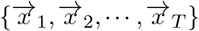 sampled from the Boltzmann distribution defined by Equation 5 for a reference state matrix 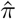. For values of *π* sufficiently close to 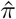 (see Methods), we define a weight

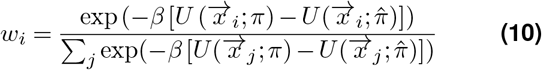

for each 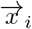 We can then express our expectation in terms of these weights

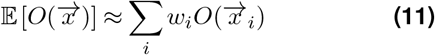

This yields an expression for 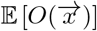 such that 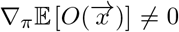. Note that 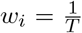 in the limit where 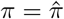. Importantly, gradients are not computed through the unrolled trajectory (as in traditional differentiable MD) but only through the energy function, relieving many of the numerical instabilities and memory constraints that typically plague differentiable MD. This is equivalent to a low-variance REINFORCE gradient estimator by using knowledge of the unnormalized steady-state probabilities to effectively integrate over all paths yielding the same equilibrium state. Additionally, the set of reference states must not be computed at every iteration (see Methods), relaxing the computational cost imposed by running large simulations.

In practice, since the rows of *π* must be normalized, one optimizes a set of logits λ ∈ ℝ*n*×*m* that are normalized in the loss function to yield *π* at each step, i.e. *π*_*i*_ = softmax(λ_*i*_). Since Equation 9 reduces to Equation 1 only when *π* is one-hot, we anneal *π* throughout the optimization by introducing a temperature term *τ* to the normalization procedure, i.e. *π*_*i*_ = softmax(λ_*i*_*/τ*). We find that a simple linear annealing scheme using *τ*_start_ = 1.0 and *τ*_end_ = 0.01 works well in most cases.

In the general case, sampling from the distribution defined by Equation 2 is intractable because there are *m*^*n*^ possible permutations of state identities. However, this calculation becomes tractable in the case of an energy function in which the total energy is expressed as the sum of pairwise energies. Consider such an energy function for a fixed set of particle identities 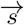:

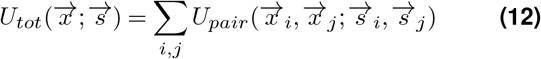

This can be generalized to the case of continuous particle identities:

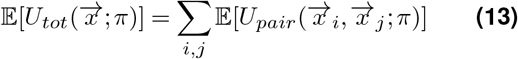

where

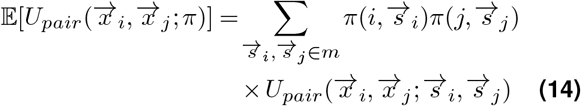

Crucially, all terms in Equation 14 are independent and we can therefore rewrite 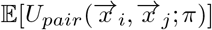 as

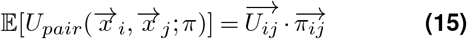

where

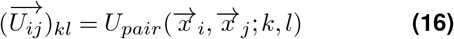

and

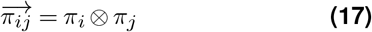

where ⊗ denotes the Kronecker product. When performed in serial, the complexity of this calculation reduces to 𝒪 (*n*^2^*m*^2^) and the *n*^2^ factor can be further reduced by the use of neighbor lists. Crucially, however, the entire calculation can be highly parallelized on a modern GPU as the terms in Equation 13 are independent. While it is standard for coarse-grained models to be pairwise, this formulation could be extended to models with *k*-body interactions where the complexity of the expected energy calculation will scale as 𝒪 (*n*^*k*^*m*^*k*^) (prior to any neighbor list optimizations).

### Differentiable Monte Carlo (DiffTRE)

Unlike a general reinforcement learning environment, we often *know things* about a physical system under study. Importantly, for example, we often know the probability distribution of the microstates of a given dynamical system. In the following, we focus on the simple case of an equilibrium system in the canonical ensemble where the probability of state 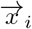 is 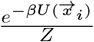 where *β* is the inverse temperature, 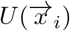 is the potential energy of 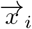, and 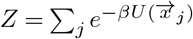 is the partition function.

Consider a set of states sampled from this distribution via some control parameters 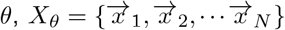 Note that there are many schemes for efficiently sampling from the Boltzmann distribution such as standard MD and Monte Carlo (MC) algorithms, and even generative deep learning methods. Examples of *θ* are parameters of the potential energy or parameters of the initial conditions (e.g. probabilities of nucleotide base identities in a simulation of nucleic acids). Via ergodicity, we can compute the expectation of some state-level observable 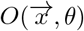 as

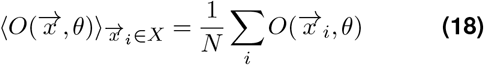

This time, our expectation is defined with respect to a set of sampled states (whose probability distribution we know) rather than with respect to a set of trajectories (or equivalently, random seeds). When formulated in this fashion, our calculation of the expectation has *no history dependence*; we do not care how the states are sampled, only that they are sampled from the underlying distribution.

However, we cannot immediately compute an accurate gradient of Equation 18. Although we know that the relative probabilities of each microstate will change as we change *θ*, we lose this dependence in our gradient signal by only considering the final set of sampled states as 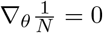 To recover this signal, Zhang et al. (29) and Thaler and Zavadlav (28) independently introduced a simple reweighting scheme (termed Differentiable Trajectory Reweighting, or DiffTRE by the latter publication) in which we rewrite Equation 18 as

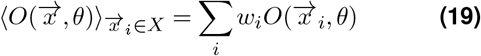

where

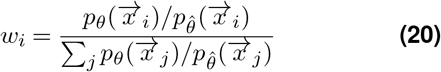

and 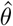 is the reference potential via which *X*_*θ*_ was sampled. Equation 20 only requires unnormalized probabilities as the normalizing factors cancel. For example, in the case of the canonical ensemble, Equation 20 does not require knowledge of the partition function:

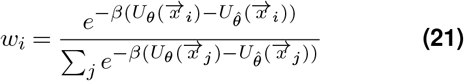

Crucially, in the case where 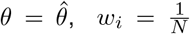 but 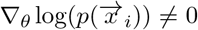 Thaler and Zavadlav introduced the notion that reference states collected via 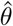 can be reused for small differences between *θ* and 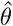, but as this difference grows few states dominate the average and the reference states should be resampled. This is captured via an expression for effective sample size:

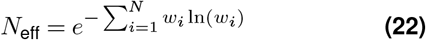

See Refs. (29) and (28) for a complete introduction to this method.

This reweighting scheme solves three major problems in differentiable programming for dynamical systems. Foremost, it resolves both problems related to memory, and numerical instability as gradients are no longer computed with respect to unrolled trajectories. However, there is a third benefit – the entire sampling procedure (e.g. simulation code) does not have to be rewritten in an automatic differentiation framework. Instead, one only must write the energy function in such a framework. In addition, objective functions that do not explicitly depend on *θ* also do not have be differentiable, permitting the immediate use of the rich ecosystem of libraries that already exist for the analysis of MD trajectories. This reduces a massive barrier to entry for differentiable programming in cases where the unnormalized probability of sampled states is known, particularly as it relates to larger and more complex code bases.

In the language of stochastic gradient estimators, DiffTRE can be regarded as a low-variance REINFORCE estimator. A traditional REINFORCE estimator would regard the probability of each state as the probability of its corresponding trajectory, drastically inflating the variance of the estimator as many trajectories can yield the same equilibrium state. DiffTRE permits us to use our knowledge about the distribution from which we are sampling in our estimate of the gradient, effectively integrating over all trajectories for a given state.

### Mpipi Force Field

Mpipi is a coarse-grained model of protein-protein and protein-RNA interactions for studying biomolecular liquidliquid phase separation (LLPS) (21). Introduced in 2021, Mpipi has gained widespread popularity for the computational study of LLPS and the underlying biophysics (55– 59). Recent machine learning methods use Mpipi to generate ground truth training data with which neural networks are trained to either predict ensemble properties or generate sequences with target characteristics (23, 60). Note that such methods for inverse design are limited not only because they generate sequences with respect to a learned approximation of Mpipi rather than Mpipi itself, but also because in principle designing sequences for a different target ensemble property demands an entirely new deep learning model.

In Mpipi, each monomer (i.e. amino acid or nucleic acid) is represented a single isotropic sphere. Each monomer type (i.e. amino acid or nucleotide identity) is assigned a mass, diameter, charge, and energy scale. Like oxDNA, all interactions are pairwise and the potential energy is given by

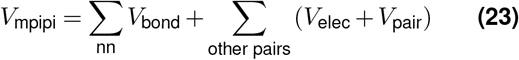

where *nn* denotes a fixed set of consecutive bonded pairs. *V*_bond_ is computed as a harmonic bond potential, *V*_elec_ as a Coulomb term with Debye-Hückel electrostatic screening, and *V*_pair_ as a Wang-Frenkel interaction (61). The parameters of this potential were fit to reproduce both atomistic potential-of-mean-force calculations and bioinformatics data. See (21) for complete details of the model and its parameterization, and see (23) for a description of the modified parameters used in this work.

### Simulations

All simulations with were performed in JAX-MD (27) on an NVIDIA A100 80 GB GPU. We used a Langevin thermostat with a timestep of 10 fs at standard conditions of 300K and 150 mM salt concentration unless specified otherwise. We make all code available via the following GitHub repository: https://github.com/rkruegs123/idp-design.

## Acknowledgments

We thank Max Ward for his collaboration on sequence design via overparameterization in the context of differentiable RNA folding, which inspired this work, Jamie Smith for helpful discussions relating to stochastic gradient estimators, and Wilton Snead for helpful discussions on IDP design. R.K.K., M.P.B., and K.S. acknowledge support from the Simons Foundation through the Simons Foundation Investigator award. R.K.K and M.P.B. acknowledge support from the NSF AI Institute of Dynamic Systems (#2112085), Office of Naval Research (N00014-17-1-3029), and the Harvard Materials Research Science and Engineering Center (DMR 20-11754). K.S. acknowledges support from NSF–Simons Center for Mathematical and Statistical Analysis of Biology at Harvard (Award #1764269) and Northwestern University for startup funding.

