## Supplementary Information for "Generalized design of sequence-ensemble-function relationships for intrinsically disordered proteins"

#### Supplementary Note 1: Stochastic Gradient Estimators for Dynamical Systems

In this work, we employ trajectory reweighting for low-variance gradient estimation through simulations. Here we provide details on traditional stochastic gradient estimation for dynamical systems.

##### A Stochastic Gradient Estimation

The problem of computing the gradient of an expectation of a function with respect to parameters defining the distribution that is integrated is well-studied in machine learning (see (62) for a complete review). Consider an objective $\mathcal{F}$ of the form

$$\mathcal{F}(\theta) := \int p(x;\theta) f(x;\theta) dx = \mathbb{E}_{p(x;\theta)}[f(x;\theta)] \tag{24}$$

One is typically concerned with finding extrema of $\mathcal{F}$, making its gradient of interest:

$$\nabla_\theta \mathcal{F}(\theta) = \nabla_\theta \left[ \int p(x;\theta) f(x;\theta) dx \right] \tag{25}$$

$$= \int p(x;\theta) \nabla_\theta f(x;\theta) dx + \int \nabla_\theta p(x;\theta) f(x;\theta) dx \tag{26}$$

$$= \int p(x;\theta) \nabla_\theta f(x;\theta) dx + \int p(x;\theta) \nabla_\theta \log(p(x;\theta)) f(x;\theta) dx \tag{27}$$

$$= \mathbb{E}_{p(x;\theta)}[\nabla_\theta f(x;\theta)] + \mathbb{E}_{p(x;\theta)}[\nabla_\theta \log(p(x;\theta)) f(x;\theta)] \tag{28}$$

In traditional supervised learning, batches are uniformly subjected to backpropagation so $\nabla \log(p(x;\theta)) = 0$ and $\nabla_\theta \mathcal{F}(\theta) = \mathbb{E}_{p(x;\theta)}[\nabla_\theta f(x;\theta)]$.

Conversely, in reinforcement learning, the distribution $p(x;\theta)$ (defined by the policy) explicitly depends on $\theta$ but the reward for a given state $f(x;\theta)$ typically does not. One solution to this is to *reparameterize* the source of the stochasticity. Under the *pathwise gradient estimator*, rather than generating samples from the distribution $p(x;\theta)$, samples are first generated from a distribution $p(\varepsilon)$ that is independent of $\theta$ and only then are these generated samples are transformed via a deterministic path $g(\epsilon, \theta)$ (i.e. a sampling process). Like the case of supervised learning, the reward now explicitly depends on $\theta$ (via the sampling process) and our distribution does not depend on $\theta$ (i.e. $\nabla_\theta p(\varepsilon) = 0$). So, the gradient calculation acts directly on the sequence of operations that are applied to sources of randomness to yield the objective function:

$$\nabla_\theta \mathcal{F}(\theta) = \mathbb{E}_{p(\varepsilon)}[\nabla_\theta f(g(\varepsilon, \theta))] \tag{29}$$

In reinforcement learning, this is referred to as the *reparameterization trick*.

There is a second, alternative gradient estimator in reinforcement learning that does not differentiate through the sampling procedure itself but only requires that the probability of sampling each state is differentiable. Traditionally, the probability of each state is represented as the joint probability of each individual step (as determined by the policy) in the trajectory that yielded a given state. Under this *score-function gradient estimator*, one deals instead with the second term in Equation 28:

$$\nabla_\theta \mathcal{F}(\theta) = \mathbb{E}_{p(x;\theta)}[\nabla_\theta \log(p(x;\theta)) f(x;\theta)] \tag{30}$$

This estimator is referred to as the REINFORCE algorithm in the reinforcement learning community. Intuitively, it provides signal to increase the probability of high reward states and decrease the probability of low reward states.

##### B Differentiable Molecular Dynamics

In traditional molecular dynamics simulations, a system of $n$ interacting bodies, typically represented by a vector $\vec{x} \in \mathcal{R}^{6n}$ representing their positions and momenta, is iteratively propagated through time via a step function $\mathcal{S}$:

$$\vec{x}_{t+1} = \mathcal{S}(\vec{x}_t, \theta)$$

where $s$ depends on the energy function and numerical integration scheme and $\theta$ are control variables. For some fixed time $N$, we can represent the final state $\vec{x}_N$ as a single function

$$\mathcal{T}(\vec{x}_0) = \mathcal{S}(\cdots \mathcal{S}(\mathcal{S}(\vec{x}_0)) \cdots) = \vec{x}_N \tag{31}$$

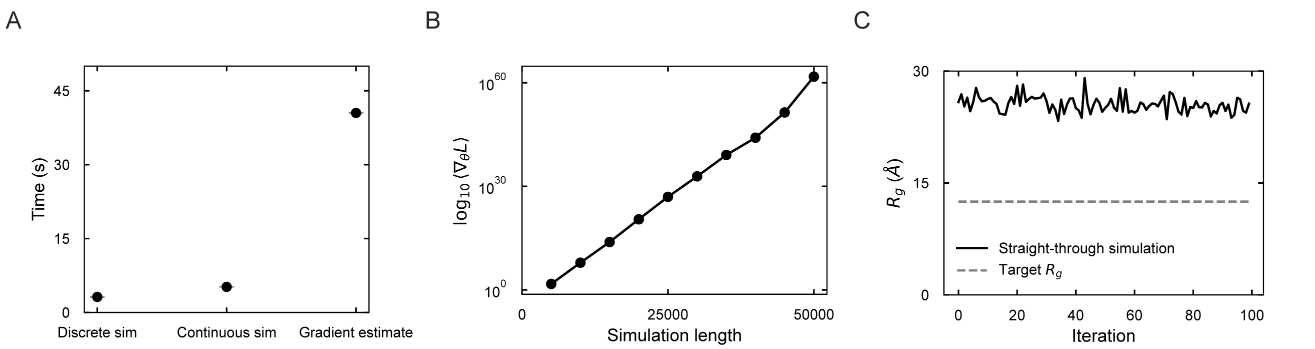


**Fig. S1. Benchmarking a differentiable implementation of the Mpipi force field**
**A.** A comparison of simulation times for a sequence of length $n = 50$ between forward simulations with discrete and continuous sequence representations, and a gradient calculation directly through the unrolled simulation with the continuous sequence.
**B.** Scaling of the mean absolute value of the gradient (on a log scale) as a function of simulation length.
**C.** An optimization trajectory using gradients computed directly through the unrolled simulation of predicted $R_g$ versus epochs.

where $\mathcal{S}$ is applied $N$ times and $\vec{x}_0$ represents the initial state. Thus, a MD trajectory can be considered the result of a single numerical calculation.

When written in an automatic differentiation framework, gradients can be computed efficiently *with respect to* this calculation. Consider some objective function defined with respect to the final state and some control parameters $\theta$, $O(\vec{x}_N, \theta)$. Since MD trajectories are stochastic, one is typically interested in the expectation of this objective function,

$$\langle O(\vec{x}_N, \theta)\rangle_{\rho\in\mathrm{R}} = \langle O(\mathcal{T}(\vec{x}_0), \theta)\rangle_{\rho\in\mathrm{R}} \quad \textbf{(32)}$$

$$= \frac{1}{|\mathrm{R}|}\sum_{\rho\in\mathrm{R}} O(\mathcal{T}_\rho(\vec{x}_0), \theta) \quad \textbf{(33)}$$

where $\mathrm{R}$ is a set of random seeds for initializing trajectories and $\vec{x}_{N,\rho}$ is the final state resulting from seed $\rho \in \mathrm{R}$. Therefore, in the language of stochastic gradient estimators, gradients are typically computed via the reparameterization trick (c.f. (63)):

$$\nabla_\theta \mathbb{E}\left[O(\vec{x}_N, \theta)\right]_{\rho\in\mathrm{R}} \approx \langle \nabla_\theta O(\mathcal{T}_\rho(\vec{x}_0), \theta)\rangle_{\rho\in\mathrm{R}} \quad \textbf{(34)}$$

Note that (i) objective functions can also be defined with respect to the entire trajectory itself rather than only the final state and (ii) in the case of equilibrium systems, one long simulation with states sampled at sufficiently long time intervals can be interpreted as a set of individual trajectories under standard assumptions of ergodicity, but we retain the current formalism for clarity.

Since gradients must be computed with respect to the simulation procedure itself, prohibitive numerical and memory limitations arise. In (64), Metz et al. derived the analytical gradient for a single term in Equation 34:

$$\frac{dO_N}{d\theta} = \frac{\partial O_N}{\partial \theta} + \sum_{k=1}^{N} \frac{\partial O_N}{\partial \vec{x}_N}\left(\prod_{i=k}^{N} \frac{\partial \vec{x}_i}{\partial \vec{x}_{i-1}}\right)\frac{\partial \vec{x}_k}{\partial \theta} \quad \textbf{(35)}$$

where $O_N$ represents the objective function evaluated at the final state. Importantly, the matrix of partial derivatives $\frac{\partial \vec{x}_i}{\partial \vec{x}_{i-1}}$ is the Jacobian of the dynamical system. Only when the magnitude of all eigenvalues of this Jacobian are less than one will the resulting product be well-behaved; otherwise, the product will diverge. Moreover, as MD trajectories often require very large numbers of steps, the memory required to compute Equation 35 can often far exceed the limitations of state of the art GPUs.

#### Supplementary Note 2: Benchmarking

Our method requires that we simulate a probabilistic sequence $\pi$ rather than a discrete sequence. For a pairwise energy function, this requires an energy calculation that is $\mathcal{O}(20^2 n^2)$ rather than $\mathcal{O}(n^2)$. In Figure S1A, we show that this incurs only a modest computational cost for a system of size $n = 50$. Given current limitations of JAX-MD, we do not use neighbor

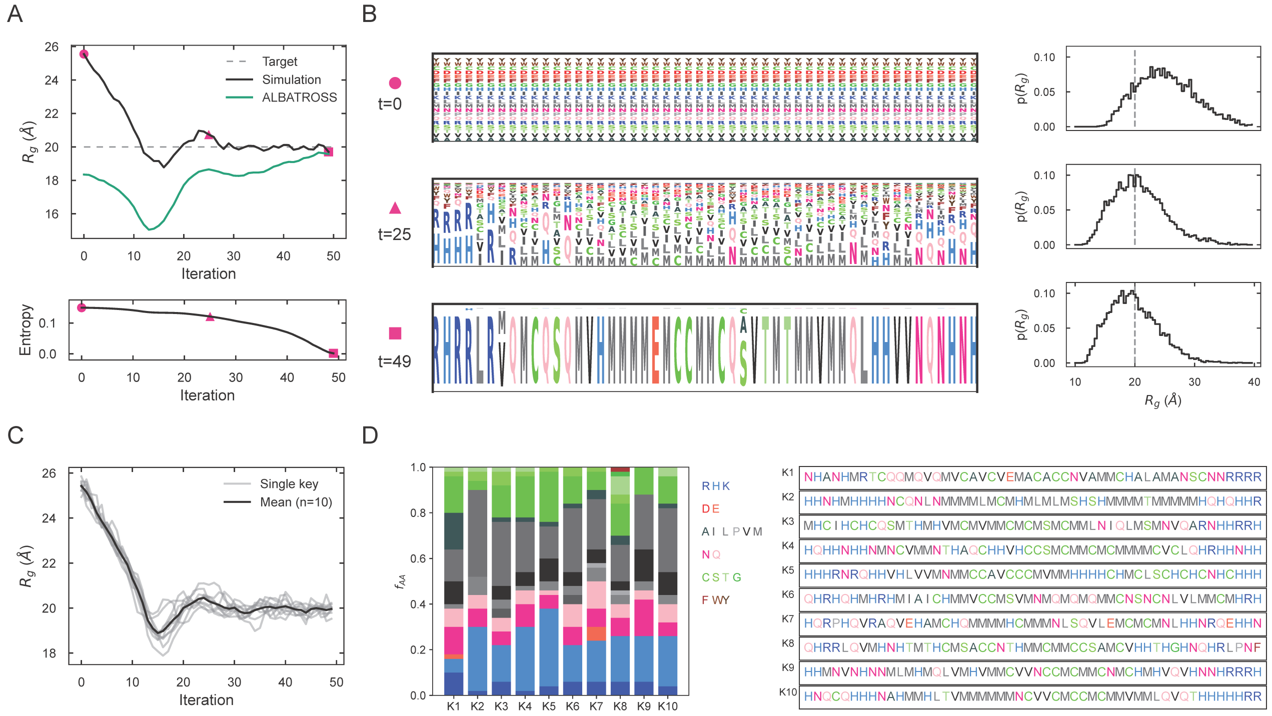


**Fig. S2. Flexibility of inverse design framework**

**A.** An explicit comparison of simulation predicted $R_g$ (in Figure 2A) to ALBATROSS, a neural network trained on the Mpipi-GG force field to predict $R_g$ from sequence, over time. The black curve represents the $R_g$ from the simulated probabilistic sequence and the green curve represents the average$R_g$ of discrete sequences sampled from the distribution of sequences defined by the probabilistic sequence as predicted by ALBATROSS. Highlighted points (in pink) represent the start, mid-point, and end of the optimization trajectory.
**B.** The evolution of the probabilistic sequence throughout the optimization depicted in (A) for each highlighted trajectory point shown through a WebLogo-style plot where height of the amino acid letter is proportional to its relative probability. On the right panel, the probability distribution of $R_g$ values across the conformational ensemble is shown.
**C.** The $R_g$ of the simulated probabilistic sequence versus epochs for ten trajectories initialized with random keys (gray lines) but with the same target $R_g = 20$ Å and the black line represents the average profile.
**D.** The left panel represents the composition of each of the 10 optimized sequences that were obtained from different random keys (x-axis) and the height of the bar is proportional to the frequency at which a particular residue appears in the sequence. To the right, the final sequences are listed.

lists in this work but the efficient usage of neighbor lists should permit the scaling of this relationship to arbitrarily long sequences.

Traditional differentiable MD requires the calculation of gradients through the unrolled trajectory. In Figure S1A, we also shows how such a gradient calculation imposes a significant time overhead to the otherwise forward simulation. Moreover, this gradient calculation is unstable as gradients explode with increasing numbers of timesteps (Figure S1B). These gradients cannot be used for optimization even in the context of a simplified version of the most basic design problem considered in this work, i.e. designing a sequence with a target $R_g$ *without* annealing the entropy of the probabilistic sequence (Figure S1C).

#### Supplementary Note 3: $R_g$ Optimizations

In Figures 2 and 4, we report sequence optimizations for target $R_g$ values. This requires a definition of $R_g$ for a probabilistic sequence. In the context of IDPs, the $R_g$ is defined as the root mean square distance of particles from the center of mass,

$$R_g^2 = \frac{1}{n}\sum_{i=1}^{n} r_i^2 \quad \textbf{(36)}$$

where $r_i^2$ denotes the squared distance of the $i^{th}$ particle to the center of mass. While the particle positions are independent of the particle identities, the center of mass is not as in general the mass of a particle depends on its residue type.

| n | Target Rg | Simulated Rg | ALBATROSS Rg |
|---|---|---|---|
| 50 | 32.5 | 32.4 | 28.5 |
| 75 | 42.5 | 41.3 | 39.0 |
| 75 | 45 | 43.4 | 37.8 |
| 75 | 47.5 | 46.1 | 39.7 |
| 75 | 50 | 48.6 | 39.0 |

**Table S1.** $R_g$ optimizations for which ALBATROSS underpredicts the simulated $R_g$.

Consider a force field that assigns masses $\overrightarrow{m}$ to each residue where $\overrightarrow{m}_j$ is the mass of the $j^{th}$ residue and $|\overrightarrow{m}| = 20$. Given a probabilistic sequence $\pi$, we define the mass of the $i^{th}$ residue as $\pi_i \cdot \overrightarrow{m}_i$ where $\cdot$ denotes the dot product. For a sequence of length $n$, we can then define the center of mass as

$$\overrightarrow{x}_{\text{COM}} = \frac{\sum_{i=1}^{n} (\pi_i \cdot \overrightarrow{m}_i) \overrightarrow{x}_i}{\sum_{i=1}^{n} \pi_i \cdot \overrightarrow{m}_i} \quad \textbf{(37)}$$

where $\overrightarrow{x}_i$ denotes the position of the $i^{th}$ particle.

Since $R_g$ is defined for a single state, the $R_g$ values reported in Figures 2 and 4 are expected values over a trajectory, i.e. $R_g^{\text{sim}} = \mathbb{E}[R_g(\overrightarrow{x})]_{\overrightarrow{x} \sim p(\cdot;\pi)}$. For optimization, we define the loss as $\mathcal{L}(\pi) = \text{RMSE}(R_g^{\text{sim}}, R_g^{\text{target}})$ where $R_g^{\text{target}}$ is the target value.

In Figure 2, we also compare the simulated $R_g$ to the average $R_g$ of discrete sequences as predicted by ALBATROSS. ALBATROSS is a model developed by Lotthammer et al. that is trained on Mpipi simulations to predict single-molecule properties such as $R_g$ and $R_{ee}$ (23). Given a probabilistic sequence $\pi \in \mathbb{R}^{n \times 20}$ (i.e. the parameter of a product of categorical distributions), discrete sequences can be sampled according to Equation 3 by independently sampling residue identities at each position. To compute $\sum_{\overrightarrow{s} \in S} p(\overrightarrow{s}|\pi) O_{\overrightarrow{s}}$ (with $O_{\overrightarrow{s}} = \mathbb{E}[O(\overrightarrow{x})]_{\overrightarrow{x} \sim p(\cdot;\overrightarrow{s})}$) via ALBATROSS, we sample 1000 discrete sequences and compute the average $R_g$ as predicted by ALBATROSS. We find that 1000 samples yields a converged average. We access ALBATROSS via the `sparrow` package made available at the following link: https://github.com/idptools/sparrow.

#### Supplementary Note 4: $R_{ee}$ Optimizations

In Figure 2, we also report sequences optimized for target values of $R_{ee}$. Unlike $R_g$, $R_{ee}$ is only a function of particle positions and therefore has the same definition for a continuous sequence as for a discrete sequence. Specifically, for a sequence of length $n$, $R_{ee}$ is defined as simply the distance between the first and last particles:

$$R_{ee} = d(\overrightarrow{x}_1, \overrightarrow{x}_n) \quad \textbf{(38)}$$

where $d$ denotes the Euclidean distance. The loss is defined similarly as in the case of $R_g$, i.e. $\mathcal{L}(\pi) = \text{RMSE}(R_{ee}^{\text{sim}}, R_{ee}^{\text{target}})$ where $R_{ee}^{\text{target}}$ is the target value and $R_{ee}^{\text{sim}} = \mathbb{E}[R_{ee}(\overrightarrow{x})]_{\overrightarrow{x} \sim p(\cdot;\pi)}$.

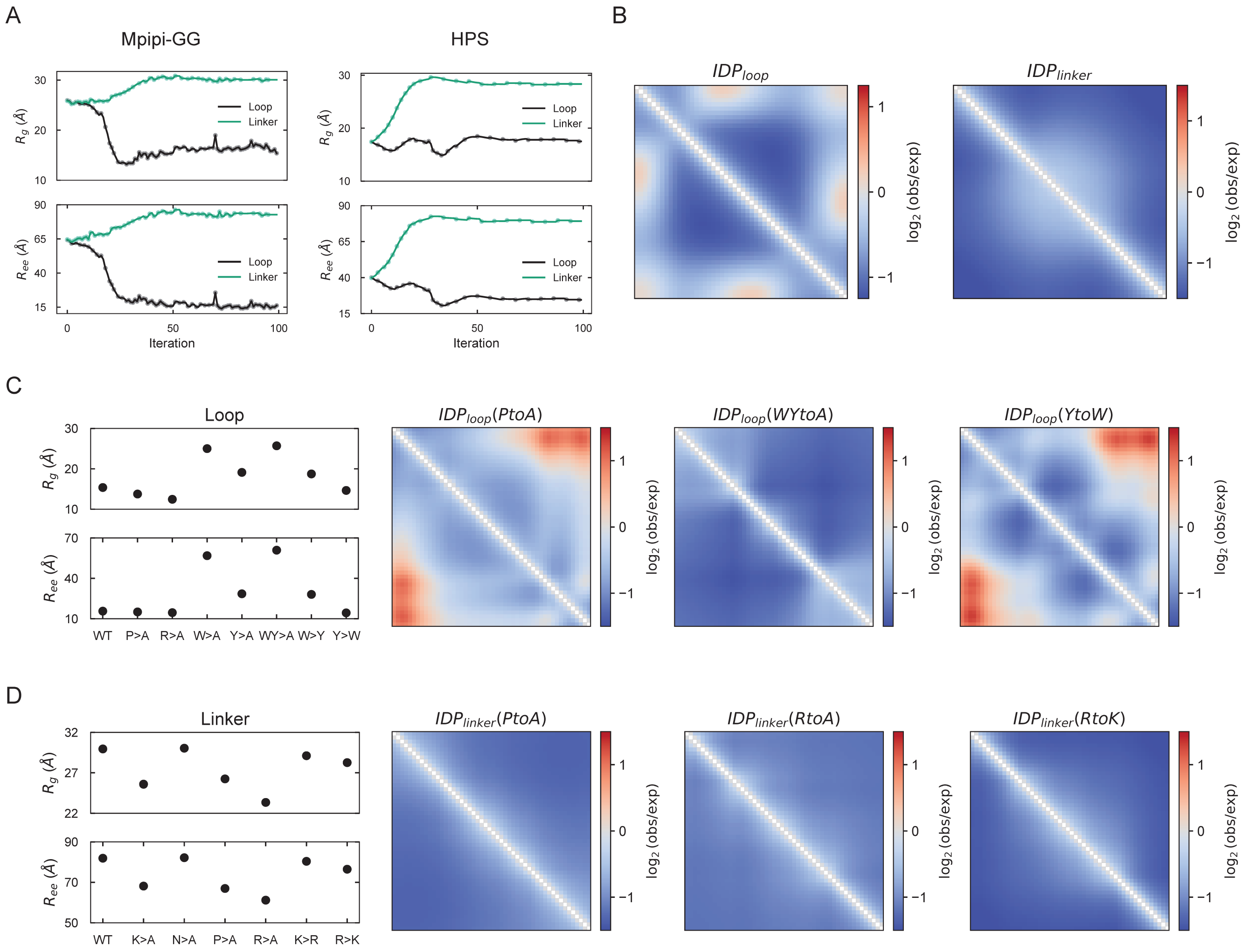


**Fig. S3. Probing mechanisms of loop and linker assembly**

**A.** Each panel depicts the convergence of $R_g$ and $R_{ee}$ over the optimization iterations for representative loop (black lines) and linker (green lines) optimizations with $n = 50$ for the Mpipi-GG (left panels) and HPS (right panels) force-fields. **B.** The panels depict normalized contact maps over representative trajectories for the loop (left) and linker (right) solutions under the HPS force field, analogous to the Mpipi-GG force field depicted in Figure 3. For contact frequencies, red/blue regions represent higher/lower expected frequencies when contrasted with an ideal polymer of identical length.

**C.** The $R_g$ (top) and $R_{ee}$ (bottom) are shown for both the optimized loop solution (WT or wild-type) and a set of mutational scans. On the right, the corresponding contact maps (as in B.) are shown for a subset of mutants.

**D.** The $R_g$ (top) and $R_{ee}$ (bottom) are shown for both the optimized linker solution (WT or wild-type) and a set of mutational scans. On the right, the corresponding contact maps (as in B.) are shown for a subset of mutants.

#### Supplementary Note 5: Loop and Linker Optimizations

In Figures 3C.I-II, we present the optimizations of loops and linkers – IDPs for which $R_g >> \frac{R_{ee}}{\sqrt{6}}$ and $\frac{R_{ee}}{\sqrt{6}} >> R_g$, respectively. For a given probabilistic sequence $\pi$, we define the loss functions as

$$\mathcal{L}_{loop}(\pi) = \frac{R_{ee}^{sim}}{\sqrt{6}} - R_g^{sim} \tag{39}$$

and

$$\mathcal{L}_{linker}(\pi) = R_g^{sim} - \frac{R_{ee}^{sim}}{\sqrt{6}} \tag{40}$$

Figure S3 depicts the convergence of the $R_g$ and $R_{ee}$ values for the representative optimizations depicted in Figures 3C.I-II. Table S2 lists the results of the mutational analyses summarized in the insets of Figures 3C.I-II.

| Sequence | $\mathbf{R_g}$ | $\mathbf{R_{ee}}$ | $\mathcal{L}oss$ |
|---|---|---|---|
| Solution | 15.4 | 15.6 | -9.0 |
| P>A | 13.8 | 15.0 | -7.6 |
| R>A | 12.5 | 14.5 | -6.5 |
| W>A | 25.0 | 56.8 | -1.8 |
| Y>A | 19.1 | 28.5 | -7.5 |
| W>Y | 18.7 | 28.1 | -7.3 |
| Y>W | 14.6 | 14.3 | -8.8 |
| Min. $R_{ee}$ | 11.7 | 13.8 | -6.1 |
| Max. $R_g$ | 36.2 | 93.9 | 2.1 |

**(a)** Loop

| Sequence | $\mathbf{R_g}$ | $\mathbf{R_{ee}}$ | $\mathcal{L}oss$ |
|---|---|---|---|
| Solution | 30.0 | 81.9 | -3.5 |
| K>A | 25.6 | 68.1 | -2.2 |
| N>A | 30.0 | 82.2 | -3.5 |
| P>A | 26.2 | 66.9 | -1.1 |
| R>A | 23.4 | 61.1 | -1.6 |
| K>R | 29.1 | 80.4 | -3.7 |
| R>K | 28.3 | 76.5 | -3.0 |
| Max. $R_{ee}$ | 37.2 | 96.6 | -2.2 |
| Min. $R_g$ | 10.6 | 17.9 | 3.3 |

**(b)** Linker

**Table S2.** Mutational analysis for the loop and linker optimizations. Loss column for loop and linker tables represents $\frac{R_{ee}}{\sqrt{6}} - R_g$ and $R_g - \frac{R_{ee}}{\sqrt{6}}$, respectively.

### Supplementary Note 6: Sequence Constraints

#### A Constraint Activation Function

Consider a function $C : \mathbb{R}^{n\times 20} \to \mathbb{R}$ that accepts a probabilistic sequence $\pi$ as input and returns a scalar value. We wish to design a sequence that satisfies a minimum value $C_{\text{min}}$ of this function. Following the work of Krueger and Ward (34), we define a ReLU function that increases sharply below the minimum value and increases slowly above this threshold:

$$\phi_C(C_\pi) = \left\{ \begin{array}{ll} m_1 C_\pi + (1 - m_1 C_{\text{min}}), & \text{for } C_\pi < C_{\text{min}} \\ m_2 C_\pi + (1 - m_2 C_{\text{min}}), & \text{for } C_\pi \geq C_{\text{min}} \end{array} \right\} \tag{41}$$

where $C_\pi = C(\pi)$ and $m_1$ and $m_2$ are hyperparameters with $m_1 << m_2 \leq 0$. Note that $\phi_C$ is defined such that $\phi_C(C_{\text{min}}) = 1.0$ and values $C_\pi < C_{\text{min}}$ are strongly penalized while values $C_\pi \geq C_{\text{min}}$ are mildly rewarded.

For a set of such constraint functions $C_i$ and their corresponding activation functions $\phi_{C_i}$, we define our objective function as the geometric mean of the baseline loss function and the activated constraints,

$$\mathcal{L}^{\text{const}}(\pi) = \mathcal{L}(\pi) \times \prod_i \phi_{C_i}(C_i(\pi)) \tag{42}$$

where $\mathcal{L}(\pi)$ is the RMSE between the simulated and target observables in the case of $R_g$ and $R_{ee}$. There are alternative methods for applying such constraints, e.g. projected gradient methods, though such methods would be complicated by our annealing of the sequence entropy and we find that the geometric mean works well in practice.

#### B Disorder Constraint

Since sequence entropy does not provide a reliable measure of disorder for probabilistic sequences, we turn to Metapredict, a machine learning model trained to predict consensus sequence disorder (35). Consider Metapredict as a function $\text{MP} : \mathbb{R}^{n\times 20} \to \mathbb{R}^n$ that maps a probabilistic sequence to an expected disorder at each position. We therefore define the disorder of a probabilistic sequence as the average predicted disorder, i.e.

$$D(\pi) = \frac{1}{n}\sum_{i=1}^{n} \text{MP}(\pi)_i \tag{43}$$

Since we use Metapredict v2, which predicts normalized measures of disorder, $D(\pi) \in [0,1]$. Note that Metapredict was only trained on discrete sequences and therefore was not intended for use with probabilistic sequences.

Though Metapredict provides a more reliable proxy for sequence disorder than sequence entropy, it still underpredicts the expected disorder for high entropy sequences and in practice we only wish to demand that the final sequence is disordered. Thus, we anneal $D_{min}$ from 0.2 to 0.8 throughout the first $80\%$ of the optimization iterations. For $\phi_D$, we define $m_1 = -1000$ and $m_2 = -0.01$.

#### C Charge Distribution Constraints

In Figure 4, we design IDP sequences with a target distribution of positively and negatively charged residues. Therefore, we require a definition for the ratio of a probabilistic sequence that is a given residue type. For clarity, we restrict attention to the case of positively charged residues.

For a discrete sequence $\overrightarrow{s}$ of length $n$, the ratio of positively charged residues is defined as

$$R_+(\overrightarrow{s}) = \frac{1}{n}\sum_{i=1}^{n}\delta_+(\overrightarrow{s}_i)$$

where $\delta_+(\overrightarrow{s}_i) = 1$ if $\overrightarrow{s}_i$ is a positively charged residue and $\delta_+(\overrightarrow{s}_i) = 0$ otherwise. Since the distribution of residues at each position is normalized in a probabilistic sequence, this definition can be generalized to such sequences:

$$\begin{aligned}
\mathbb{E}[R_+(\overrightarrow{s})]_{\overrightarrow{s}\sim\pi} &= \sum_{\overrightarrow{s}\in S} p(\overrightarrow{s}|\pi)R_+(\overrightarrow{s}) \\
&= \sum_{\overrightarrow{s}\in S} p(\overrightarrow{s}|\pi)\left(\frac{1}{n}\sum_{i=1}^{n}\delta_+(\overrightarrow{s}_i)\right) \\
&= \frac{1}{n}\sum_{i=1}^{n}\sum_{\overrightarrow{s}\in S} p(\overrightarrow{s}|\pi)\delta_+(\overrightarrow{s}_i) \\
&= \frac{1}{n}\sum_{i=1}^{n}\sum_{j=1}^{20}\pi_{ij}\delta_+(\overrightarrow{s}_i) \\
&= \frac{1}{n}\sum_{i=1}^{n}\pi_i\cdot\overrightarrow{\delta_+}
\end{aligned}$$

where $\overrightarrow{\delta_+} \in \mathbb{R}^{20}$ is a one-hot vector denoting whether or not a given amino acid is positively charged and $\cdot$ denotes the dot product.

As for disorder constraints, we apply an activation function $\phi_R$ to the computed ratios. For the ratios of both positively and negatively charged residues, we set $R_{min} = 0.495$ to relax the search space, and define $m_1 = -1000$ and $m_2 = -0.01$. In practice all presented sequences satisfy the constraints within $5\%$ error. We initialize logits corresponding to a value of $\pi$ such that each position has a $0.495$ probability of being both positively and negatively charged, and uncharged residue types uniformly distributed over the remaining cumulative probability $1.0 - 2\times 0.495 = 0.01$.

| Sensor Type | Response Type | $\mathbf{R_g^{lo}}$ | $\mathbf{R_g^{hi}}$ | $\mathcal{L}$oss |
|---|---|---|---|---|
| Salt | Contractor | 23.1 | 14.1 | -9.0 |
| Salt | Expander | 14.4 | 26.6 | -12.2 |
| Phosphorylation | Contractor | 24.4 | 23.3 | -1.1 |
| Phosphorylation | Expander | 16.7 | 19.4 | -2.7 |
| Temperature | Contractor | 32.0 | 31.3 | 0.7 |
| Temperature | Expander | 14.2 | 19.7 | -5.5 |

**Table S3.** Optimized sensors of length $n = 50$ for a range of sensor and response types. For a salt sensor, $R_g^{lo}$ and $R_g^{hi}$ correspond to 150 mM and 450 mM, respectively. For a phosphorylation sensor, $R_g^{lo}$ and $R_g^{hi}$ correspond to the $25^{th}$ position fixed as serine (S) and Glutamic acid (E), respectively. For a temperature sensor, $R_g^{lo}$ and $R_g^{hi}$ correspond to 293.15 K (20 C) and 363.15 K (90 C), respectively.

#### D Overparameterization

In Figure 4, we overparameterize the search space by optimizing over the weights of a neural network that outputs a $n\times 20$ matrix of logits. We use a fully-connected architecture for all networks with 6 layers of 4000 nodes each. We apply a Leaky ReLU activation function bewteen each layer. We pretrain the network to output a target set of logits corresponding to a target probabilistic sequence (i.e. a uniform-distributed probabilistic sequence or one with a target distribution of charged residues) Given a target initial pseq $\pi^{init}$, we define the following pretrain loss:

$$\mathcal{L}_{pretrain}(\theta) = \frac{1}{4n}\sum_{ij}100\cdot(\pi_{ij}^{pred} - \pi_{ij}^{init})^2 \tag{44}$$

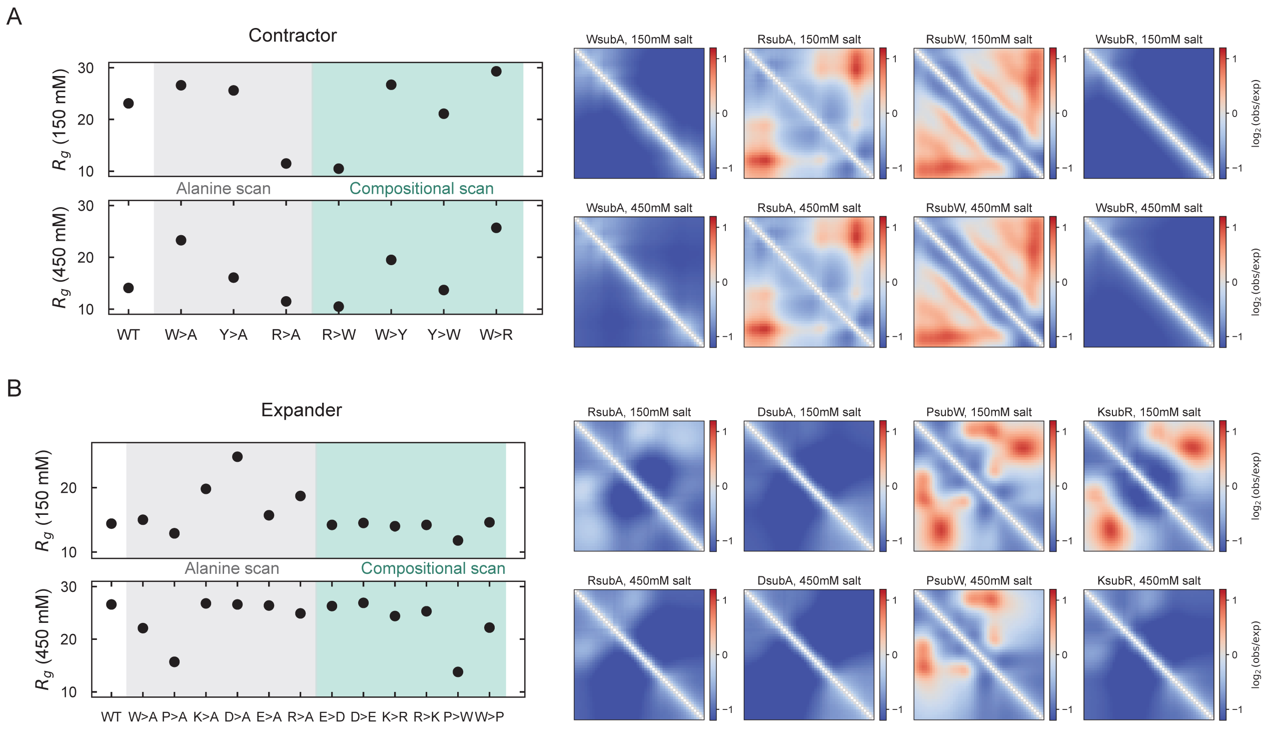


**Fig. S4. Mutational analyses for the optimized salt sensors**
**A-B.** For the salt sensing contractor (A) and expander (B) reported in Figure 5, we perform alanine scanning (gray background) as well as rational compositional mutations (green background) informed by the underlying sensor mechanism. For each of the sensors, we report the effect of particular mutations on $R_g$ at low (top subpanel) and high salt (bottom subpanel). On the right, normalized contact frequencies are shown for particular mutants for low (top subpanel) and high salt (bottom subpanel). For contact frequencies, red/blue regions represent higher/lower expected frequencies when contrasted with an ideal polymer of identical length.

where $\pi^{pred} =$ softmax(NN$_\theta(k)$), $\theta$ are the weights of the neural network, and NN$_\theta(k)$ denotes the output of the neural network with weights $\theta$ and a fixed random seed as input. We use an Adam optimizer with a learning rate of learning rate of $10^{-5}$ for pretraining.

#### Supplementary Note 7: Sensor Optimizations

##### A Salt Sensors

In Figure 5, we design IDPs that expand or contract upon the addition of salt. In Mpipi, a single Debye length $\kappa$ was used to reproduce behavior of IDPs at a salt concentration of 150 mM. To model the effects of increased salt, we followed Debye-Huckel theory in which

$$\kappa^{-1} = \sqrt{\frac{\varepsilon_R \varepsilon_0 kT}{2e^2 I}} \tag{45}$$

where I is the ionic strength, $\varepsilon_0$ is the permittivity of free space, $\varepsilon_R$ is the dielectric constant, and $e$ is the elementary charge. Thus, given the default Debye length in Mpipi $\kappa^{150}$, we obtain the Debye length for an arbitrary salt concentration $I$ (expressed in mM) as

$$\kappa^I = \kappa^{150} \frac{\sqrt{I/1000}}{\sqrt{150/1000}} \tag{46}$$

More generally, salt concentration can affect simulation parameters in multiple ways (e.g. the dielectric permittivity) and such effects can be easily accommodated in our framework through empirical models.

We formulate the sensor design problem similar to that for the design of loops and linkers, except that the loss function involves expectations from two distinct ensembles. We define two salt concentrations $I^{lo}$ and $I^{hi}$. corresponding to Debye lengths $\kappa^{lo}$ and $\kappa^{hi}$. For a given probabilistic sequence $\pi$, we compute the $R_g$ in each ensemble, denoted $R_g^{lo}$ and $R_g^{hi}$,

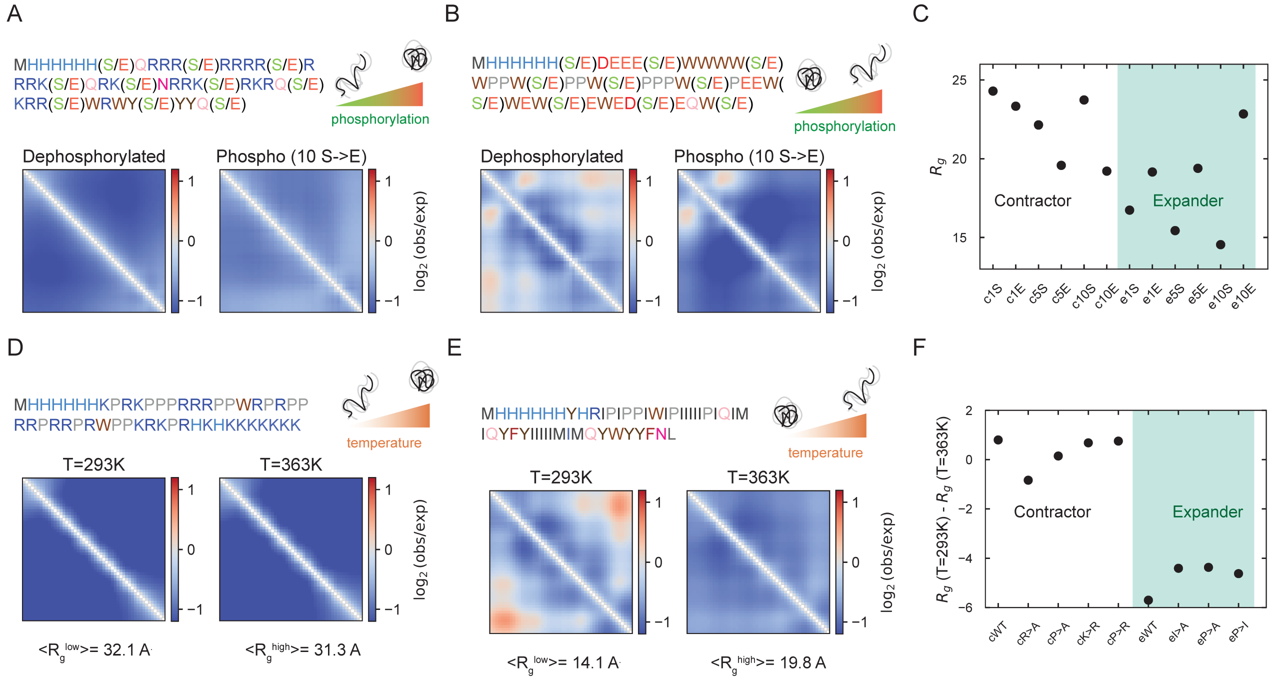


**Fig. S5. Phosphorylation and temperature sensing IDPs**
**A-B.** Optimized contractor and expander sequences with normalized contact frequency maps without (A) and with (B) phosphorylation computed from representative trajectories. For these solutions, the IDPs comprise 10 phosphosites i.e., Serines that are roughly equi-spaced, and highlighted in the sequence. For contact frequencies, red/blue regions represent higher/lower expected frequencies when contrasted with an ideal polymer of identical length.
**C.** The change in $R_g$ for optimized phosphorylation sensors that exploit different numbers of phosphosites for contractors (white background) and expanders (green background). The magnitude of the effect is generically larger with more phosphosites - the axis label denotes the type of sensor (c/e), the number of serines(1/5/10), and the phosphorylation status (S/E).
**D-E.** Optimized contractor and expander sequences with normalized contact frequency maps at low (D) and high (E) temperatures. For contact frequencies, red/blue regions represent higher/lower expected frequencies than an ideal polymer of identical length.
**F.** The change in $R_g$ for optimized temperature sensors for contractors (white background) and expanders (green background) upon mutation of key residues. The axis label denotes the type of mutant.

and define the loss for the contractor as

$$\mathcal{L}_{contractor}(\pi) = R_g^{hi} - R_g^{lo} \tag{47}$$

and the loss for the expander as

$$\mathcal{L}_{expander}(\pi) = R_g^{lo} - R_g^{hi} \tag{48}$$

As a demonstration of the flexibility of our method, we impose the constraint that the sequence begins with a M-His6 motif. This represents an experimentally-relevant constraint as the start codon is required for expression and the His6 tag is commonly used for protein purification. We impose this constraint by setting the first seven rows of the probabilistic sequence (computed via $\pi = \text{softmax}(\lambda/\tau)$ where $\lambda$ are logits and $\tau$ is a temperature parameter) to a $n \times 7$ one-hot sequence representing this motif.

We optimize contractors and expanders for lengths $n = 50$, $n = 75$, and $n = 100$. For $n = 50$, we also conduct a mutational analysis of our solutions to evaluate both the effect of individual residue types via alanine substitutions, and the Pareto front via substitution of residues with residue types similar in their electrostatic properties.

#### B Phosphorylation Sensors

We next sought to design IDPs that contract or expand upon phosphorylation rather than in the presence of salt. Since phosphorylation is not explicitly modelled in Mpipi, we choose a residue pair representative of a phosphorylation event and modelled phosphorylation as the transition from one residue to the other; for our purposes, we considered the

| | Sequence | $\mathbf{R_g^{lo}}$ | $\mathbf{R_g^{hi}}$ | $\mathcal{L}$oss |
|---|---|---|---|---|
| | Solution | 23.1 | 14.1 | -9.0 |
| Alanine Scanning | W>A | 26.6 | 23.3 | -3.3 |
| | Y>A | 25.6 | 16.1 | -9.5 |
| | R>A | 11.5 | 11.5 | 0.0 |
| Conformational Scanning | R>W | 10.5 | 10.5 | 0.0 |
| | W>Y | 26.7 | 19.5 | -7.2 |
| | Y>W | 21.1 | 13.7 | -7.4 |
| | W>R | 29.3 | 25.7 | -3.6 |
| | $R_{50}$ | 30.4 | 26.8 | -3.6 |

**(a)** Contractor

| | Sequence | $\mathbf{R_g^{lo}}$ | $\mathbf{R_g^{hi}}$ | $\mathcal{L}$oss |
|---|---|---|---|---|
| | Solution | 14.4 | 26.6 | -12.2 |
| Alanine Scanning | W>A | 15.0 | 22.1 | -7.1 |
| | P>A | 12.9 | 15.7 | -2.8 |
| | K>A | 19.8 | 26.8 | -7.0 |
| | D>A | 24.8 | 26.6 | -1.8 |
| | E>A | 15.7 | 26.4 | -10.7 |
| | R>A | 18.7 | 24.9 | -6.2 |
| Conformational Scanning | E>D | 14.2 | 26.3 | -12.1 |
| | D>E | 14.5 | 26.9 | -12.4 |
| | K>R | 14.0 | 24.4 | -10.0 |
| | R>K | 14.2 | 25.3 | -11.1 |
| | P>W | 11.8 | 13.8 | -2.0 |
| | W>P | 14.6 | 22.2 | -7.6 |
| | $E_{25}K_{25}$ | 13.4 | 15.6 | -2.2 |

**(b)** Expander

**Table S4.** Mutational analysis via alanine substitutions and rational substitutions for the optimized salt sensors depicted in Figure 5. Loss column for contractor and expander tables represents $R_g^{hi} - R_g^{lo}$ and $R_g^{lo} - R_g^{hi}$, respectively.

phosphorylation of serine (S) to Glutamic acid (E). For a given length $n$, we choose a phosphorylation position $1 \leq i_{phos} \leq n$ and model the dephosphorylated and phosphorylated ensembles by explicitly setting the $i_{phos}^{th}$ residue to be (a one-hot vector representing) S and E, respectively. The loss function is defined as in Equations 47 and 48 where $R_g^{lo}$ and $R_g^{hi}$ correspond to the dephosphorylated and phosphorylated ensembles, respectively, and we similarly impose a M-His6 prefix.

#### C Temperature Sensors

We also design IDPs that contract or expand in response to changes in temperature. The optimization problem is defined as above with $R_g^{lo}$ and $R_g^{hi}$ corresponding to 293.15 K (20 C) and 363.15 K (90 C), respectively, and we again impose a M-His6 prefix. For simplicity, we do not modify $\kappa$ in accordance with the change in $kT$.

### Supplementary Note 8: Binder Optimizations

In Figure 6, we present the optimization of binder sequences for a fixed substrate. We use a simple formulation of the binder design problem in which we optimize for the binder sequence that minimizes the interstrand distance. The interstrand distance $r_{com}$ is defined as the distance between the center of masses $\vec{x}_{\text{com}}^{binder}$ and $\vec{x}_{\text{com}}^{substrate}$, each defined following Equation 37. To maximize the probability of sampling configurations with low interstrand distance, we apply a bias potential

$$U^{bias}(r_{com}) = \begin{Bmatrix} k(r_{com} - r_{max})^2, & \text{if } r_{com} > r_{max} \\ 0, & \text{otherwise} \end{Bmatrix} \tag{49}$$

To correct this bias, we redefine the probability of a state in DiffTRE ($p_\theta(\vec{x}_i)$ in Equation 20) following the standard umbrella sampling correction:

$$p_\theta(\vec{x}_i) = \frac{1}{\omega_i} \exp\left(-\beta\left(U_\theta\left(\vec{x}_i\right)\right)\right) \tag{50}$$

where $\omega_i = \exp(-\beta(U_\theta^{bias}(\vec{x}_i)))$. For the optimization depicted in Figure 6, we design a binder of length $n = 50$ and set $k = 1$ and $r_{max} = 300$.

To demonstrate the flexibility of our method, we optimize binders for two additional sequences: (1) a positively charged homopolymer (polyR) and (2) the low complexity region of Whi3 (see Figure S6). For PolyR, we design a binder of length $n = 30$ and set $k = 10$ and $r_{max} = 150$. For Whi3-LC, we design a binder of length $n = 50$ and set $k = 1$ and $r_{max} = 250$.

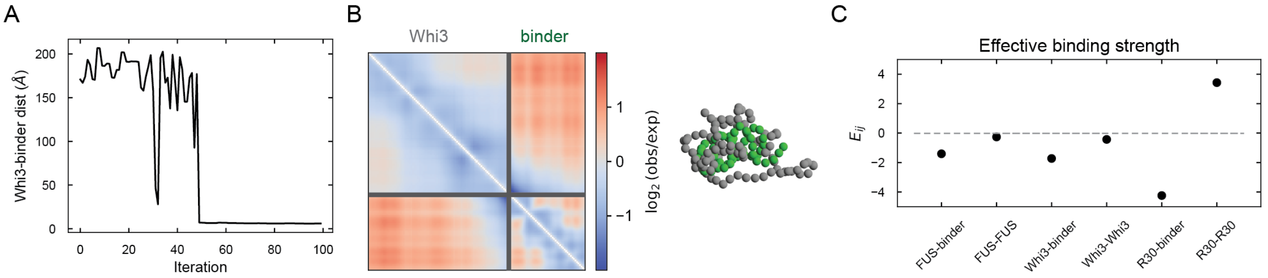


**Fig. S6. Probing IDP binder-substrate interactions**

**A.** The optimization of a binder ($n = 50$) for Whi3 ($n = 93$) is depicted by convergence to low interstrand distance over the trajectory.

**B.** Normalized contact frequency map for the optimized Whi3 binder, highlighting both intramolecular and intermolecular interactions. Red/blue regions represent higher/lower expected frequencies when contrasted with an ideal polymer of total length binder + substrate. On the right, a representative bound snapshot of binder (green) and substrate (grey) is depicted.

**C.** Computed effective interaction coefficients ($E_{ij}$, units of $nm^3$) between species $i, j$ are plotted comparing optimized binder-substrate interactions with substrate-substrate interactions. More negative values represent stronger interactions.

We then compute effective interaction coefficients to estimate the strength of interactions between substrates and ligands, as well as homotypic substrate interactions. As reported in (41), and subsequently in a related paper (42), such coefficients broadly correlate with experimental or simulation derived interaction coefficients and multicomponent condensation. Specifically, we compute the pairwise dimer coefficient ($B_{ij}^{rdp}$ as defined in (41)) between two biomolecular species $i$ and $j$, and report normalized interactions $E_{ij} = \frac{B_{ij}}{n_i n_j}$ in units of $nm^3$.
